# An Open Microfluidic Coculture Model of Fibroblasts and Eosinophils to Investigate Mechanisms of Airway Inflammation

**DOI:** 10.1101/2022.07.11.499658

**Authors:** Yuting Zeng, Xiaojing Su, Meg G. Takezawa, Paul S. Fichtinger, Ulri N. Lee, Jeffery W. Pippin, Stuart J. Shankland, Fang Yun Lim, Loren C. Denlinger, Nizar N. Jarjour, Sameer K. Mathur, Nathan Sandbo, Erwin Berthier, Stephane Esnault, Ksenija Bernau, Ashleigh B. Theberge

## Abstract

Interactions between fibroblasts and immune cells play an important role in tissue inflammation. Previous studies have found that eosinophils activated with interleukin-3 (IL-3) degranulate on aggregated immunoglobulin G (IgG) and release mediators that activate fibroblasts in the lung. However, these studies were done with eosinophil-conditioned media that have the capacity to investigate only one-way signaling from eosinophils to fibroblasts. Here, we demonstrate a coculture model of primary normal human lung fibroblasts (HLFs) and human blood eosinophils from patients with allergy and asthma using an open microfluidic coculture device. In our device, the two types of cells can communicate via two-way soluble factor signaling in the shared media while being physically separated by a half wall. Initially, we assessed the level of eosinophil degranulation by their release of eosinophil-derived neurotoxin (EDN). Next, we analyzed the inflammation-associated genes and soluble factors using reverse transcription quantitative polymerase chain reaction (RT-qPCR) and multiplex immunoassays, respectively. Our results suggest an induction of a proinflammatory fibroblast phenotype of HLFs following the coculture with degranulating eosinophils, validating our previous findings. Additionally, we present a new result that indicate potential impacts of activated HLFs back on eosinophils. This open microfluidic coculture platform provides unique opportunities to investigate the intercellular signaling between the two cell types and their roles in airway inflammation and remodeling.

## 1. Introduction

Airway inflammation is associated with a number of pulmonary diseases, including asthma (Louis et al., 2000; Angelis et al., 2014). In particular, eosinophilic airway inflammation is found in up to 80% of people with asthma and 40% with chronic obstructive pulmonary disease, indicated by an increased eosinophil count in their sputum (Pavord, 2013). Eosinophils are immune cells that play an essential role in tissue inflammation and remodeling (Lee et al., 2010). These cells can be activated to undergo degranulation and cytolysis, releasing cytotoxic granule proteins and a variety of proinflammatory soluble mediators that impact surrounding cells, including fibroblasts (Acharya and Ackerman, 2014; Spencer et al., 2014; Esnault et al., 2020). Fibroblasts are mesenchymal cells often found in connective tissues that provide structural support in various organs (Plikus et al., 2021). They play a key role in the immune response and wound healing via the uptake and secretion of various inflammatory signals (Desjardins-Park et al., 2018; Davidson et al., 2021).

In asthma, chronic airway inflammation is accompanied by the activation and differentiation of lung fibroblasts, which eventually leads to enhanced subepithelial thickness, fibrosis, and irreversible airway obstruction (Al-Muhsen et al., 2011; Torr et al., 2015; Mostaço-Guidolin et al., 2019). Eosinophils release cytokines that promote fibroblast activation, including transforming growth factor β (TGF-β) and interleukin-1 β (IL-1β) (Hinz et al., 2007; Esnault et al., 2012; Mia et al., 2014; McBrien and Menzies-Gow, 2017). Previous work has shown that eosinophil-mediated signaling could lead to either pro-inflammatory or pro-fibrotic fibroblast phenotypes (Phipps et al., 2002; Bernau et al., 2018). However, these studies did not employ *ex vivo* coculture of primary human eosinophils with primary human lung fibroblasts (HLFs), which may exhibit important differential signaling processes compared to these previous studies. Thus, coculture methods of these primary cells provide a unique advantage in the investigation of the potential mechanisms of airway inflammation via fibroblast activation.

We previously reported an *ex vivo* eosinophil degranulation model, where primary human blood eosinophils activated with interleukin-3 (IL-3) can robustly degranulate and lyse on well plates coated with heat aggregated immunoglobulin G (HA-IgG) (Esnault et al., 2017b). To study the impact of the degranulation products and cellular content from eosinophils on HLFs, we previously isolated supernatants from eosinophils in that model, and exposed them to primary lung fibroblasts, including HLFs and human bronchial fibroblasts (HBFs) (Esnault et al., 2017a; Bernau et al., 2018, 2021). In these previous studies, we observed a proinflammatory fibroblast phenotype after the treatment with eosinophil supernatants (Esnault et al., 2017a; Bernau et al., 2018, 2021). However, these studies were performed using eosinophil-conditioned media collected from these cells after they had degranulated and lysed, which only allowed one-way signaling from eosinophils to fibroblasts. This approach lacked signaling from fibroblasts to eosinophils, as well as the real-time assessment of cell-cell soluble factor bidirectional signaling.

Microfluidic coculture offers several advantages for studying cell-cell signaling over conventional approaches, such as conditioned media or Transwell^®^ inserts (Bhatia and Ingber, 2014; Li et al., 2016). Besides its ability to realize bidirectional communication in coculture, the versatile configurations of microfluidic coculture platforms provide more accurate recapitulation of the microenvironment (Young and Beebe, 2010); the flexibility in designs also enables different cell seeding ratios, culture chamber dimensions and numbers with a potential for triculture. Additionally, microfluidic devices operate with less volume, which minimizes the use of expensive reagents and conserves precious primary samples isolated from patients (Sackmann et al., 2014). Moreover, the emerging open microfluidic technology provides easy accessibility and can be operated with a pipette in a standard biological laboratory without specialized equipment (Casavant et al., 2013; Berry et al., 2017; Álvarez-García et al., 2018; Humayun et al., 2018; Berthier et al., 2019; Lee et al., 2019). In light of this, we recently developed a novel open microfluidic coculture device using the common cell culture material polystyrene, where two culture chambers are separated by a half wall, the connection of which can be temporally controlled to allow soluble factor signaling via a liquid bridge (Zhang et al., 2020) (**Figure 1**).

**FIGURE 1.**
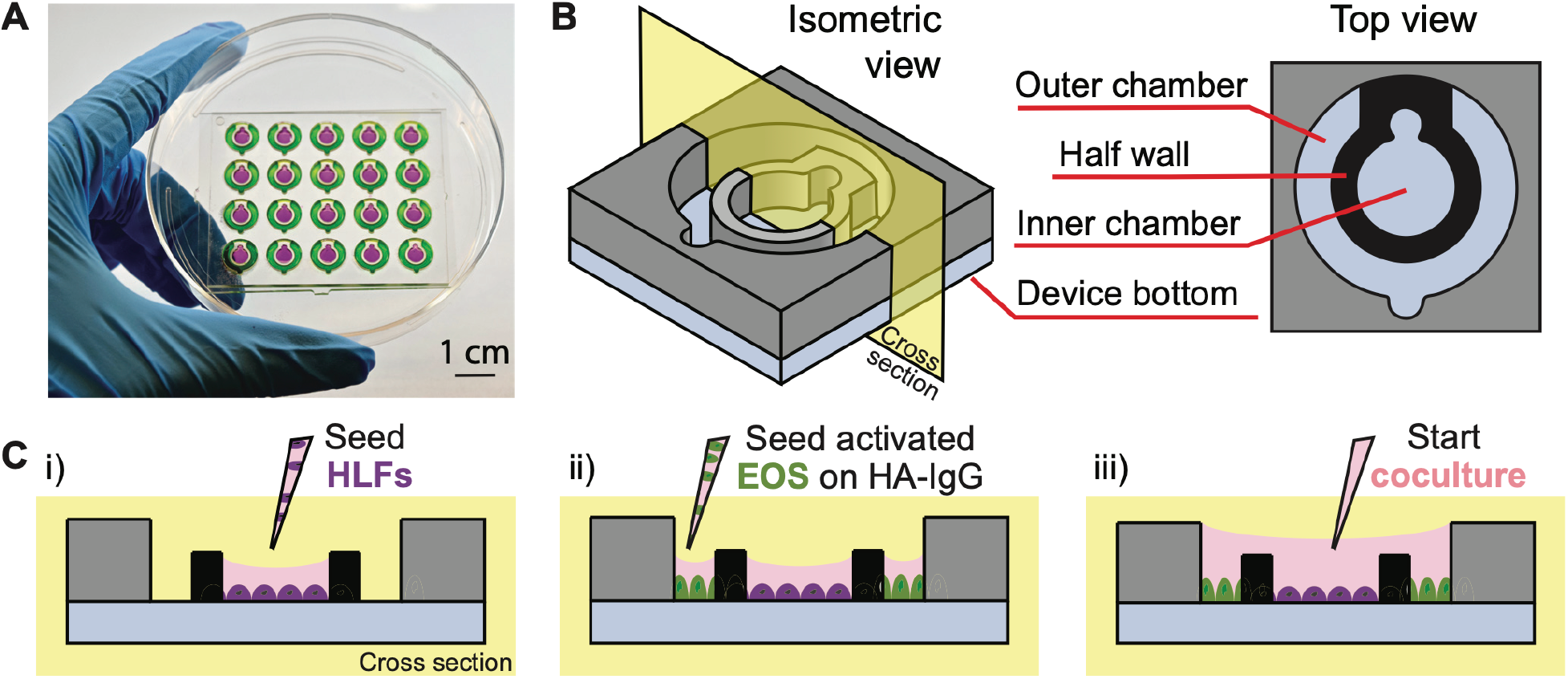
Open microfluidic coculture device description. **(A)** A device chip containing 20 coculture devices in a 4 × 5 array fitted in a 10-cm petri dish (Zhang et al., 2020). Reproduced from Zhang et al. 2020 as allowed per journal policy. Inner (purple) and outer (green) chambers are loaded with food coloring dyes for visualization. **(B)** An isometric and top view of the device design. **(C)** A cross-sectional view of the device and simplified workflow. (**i**) HLFs are seeded to the inner chambers on day 1; (**ii**) interleukin-3 (IL-3)-activated eosinophils (EOS) are seeded to the outer chambers precoated with heat-aggregated immunoglobulin G (HA-IgG) on day 3; (**iii**) inner and outer chambers are connected via the addition of coculture medium after 1.5 h to initiate coculture.

The current study builds on our prior work using this device to study cell signaling in the kidney (Zhang et al., 2020); and extends its application to include immune cell-fibroblast interactions, thus, establishing a new biological model. Further, in this work we expand the readouts possible with this device beyond fluorescence microscopy and gene expression reported in Zhang et al. 2020, demonstrating the ability to conduct phase contrast microscopy to study eosinophil morphology changes upon degranulation/cytolysis and monitor HLFs in real time (facilitated by both culture chambers being in the same focal plane). Further, the open nature of our microfluidic device (in contrast to conventional closed channels) allows for easy sampling and measurement of soluble factors using multiplexed immunoassays.

In this study, we adapt the open microfluidic coculture platform to establish a novel coculture model of degranulating eosinophils and primary HLFs to investigate the mechanisms of airway inflammation. We validate the model by characterizing eosinophil degranulation (release of eosinophil-derived neurotoxin (EDN)), HLF proinflammatory gene expression (interleukin 6 (*IL6*), C-X-C motif chemokine ligand 8 (*CXCL8*), and intercellular adhesion molecule 1 (*ICAM1*)), as well as soluble factors released in the culture media (interleukin-6 (IL-6) and interleukin-8 (IL-8)) that were previously found in the eosinophil-conditioned media studies (Esnault et al., 2017a; Bernau et al., 2018, 2021). Additionally, we present a new analyte (granulocyte-macrophage colony-stimulating factor (GM-CSF)) found in our coculture system which has not been characterized by our group before. In the future, we will continue using this validated model to discover novel bidirectional signals and the consequences on the biology of both cell types that are conferred in cocultures. Although the present study uses eosinophils derived from participants with allergy and asthma, in the future, we envision that this coculture model system could be more broadly applied to other immune cells and organs (*e*.*g*., liver, breast, prostate, gastrointestinal tract). Taken together, we demonstrate that our open microfluidic coculture device can be effectively used to dissect soluble factor signaling mechanisms between immune cells and fibroblasts.

## 2. Materials and Methods

### 2.1 Device fabrication

The open microfluidic coculture device (**Figure 1**) was designed with Solidworks (Solidworks, Waltham, MA). An engineering drawing of the device with dimensions and original design files are included in **Supplementary Figure 1** and the Supplemental Materials of the previous work (Zhang et al., 2020). The coculture devices were fabricated using a DATRONneo (Datron Dynamics, Milford, NH) Computer Numerical Control (CNC) mill. Device top and bottom pieces were milled from 2-mm and 1.2-mm thick polystyrene (PS) sheets (Goodfellow USA, Coraopolis, PA), respectively. After milling, the pieces were cleaned thoroughly with pressurized air to remove most of the plastic particles. The top and bottom pieces were then further cleaned via sonication and solvent bonded together to form a complete device chip using previously established protocols (Young et al., 2013; Zhang et al., 2020) with a few modifications to improve the bonding efficiency as described in the following paragraphs.

Modifications to the solvent bonding procedure to improve on prior protocol in Zhang et al. 2020: The inner wall of the coculture device is prone to faulty solvent bonding because it is not flush with the top surface, thus, when the weight is added the inner wall is not pressed firmly to the bottom layer for solvent bonding. A PDMS layer was designed to apply downward pressure to the inner wall during the solvent bonding process and ensure its bonding. A mold was milled from 4.5 mm poly(methyl methacrylate) on a DATRONneo. PDMS (Sylgard™ 184, Dow) base and curing agent was mixed in a 8:1 ratio and degassed in a vacuum chamber. The mixture was poured over the mold in an even layer. The mold with PDMS was degassed and cured at room temperature for 24 h. The PDMS piece, along with the 2-mm and 1.2-mm milled PS pieces, were then sonicated with 70% ethanol for 30 minutes, then was dried with pressurized air prior to solvent bonding.

For solvent bonding, a hot plate (HP60, Torrey Pines Scientific) was preheated to 65 °C. The sonicated PS and PDMS pieces and an aluminum bar (McMaster-Carr, Los Angeles, CA) were placed on the hot plate to completely dry off excess ethanol from sonication and heat each item. Once the preheating was complete, a device bottom piece was placed on a Kimwipe (Kimberly-Clark Professional™), and acetonitrile (ACN; A998-4, Fisher Chemicals) was pipetted dropwise onto the bottom piece until it covered most of the surface. The top piece was tilted at an angle such that its top edge was in contact with the longer edge of the bottom piece and was gently lowered onto the bottom piece to align the two pieces. The Kimwipe previously placed underneath the device was removed and used to gently wipe around the newly bonded device to absorb excess ACN. The heated PDMS piece was then stacked onto the device and pressure was applied to the stack with the palm of a gloved hand for about 1 min; more Kimwipes were used to absorb any excess ACN in the process. While leaving the PDMS stacked onto the device, a heated aluminum bar was placed on top to continue applying pressure for 10 min at 65 °C to facilitate the reaction between ACN and the PS interface.

After bonding, the device was sonicated again with 70% ethanol for 30 min then dried completely with pressurized air. Before cell culture use, the device was plasma treated for 5 min at 0.25 mbar and 70 W in a Zepto LC PC Plasma Treater (Diener Electronic, Ebhausen, Germany) using oxygen followed by 10 min of ultraviolet sterilization.

### 2.2 Human subjects and cell preparations

Normal HLFs were isolated as described previously (Sandbo et al., 2013; Esnault et al., 2017a) using deidentified tissue samples from thoracic surgical resection specimens. These samples were collected and characterized by the Carbone Cancer Center Translational Science BioCore at the University of Wisconsin-Madison, under Institutional Review Board approval (IRB #2016-

0934) and fibroblast subsequently derived under IRB #2011-0521. To obtain non-fibrotic fibroblasts, we utilized adjacent (uninvolved) lung from lobectomy or biopsy specimens from patients undergoing lung resection for pulmonary nodules and who did not have any identifiable lung disease by history or histologic assessment. All specimens used for fibroblast isolation were examined by a pathologist to ensure that no underlying lung disease (emphysema, idiopathic lung diseases, etc.) was present in histology. HLF cultures were derived from two different donors.

As previously, to isolate fibroblasts, tissue specimens were placed in fibroblast growth media (Dulbecco’s Modified Eagle’s Medium (DMEM; Corning 10-013-CV) with 100 units/mL streptomycin, 250 ng/mL amphotericin B, 100 units/mL penicillin (Corning 30-004-CI), and 10 μg/mL ciprofloxacin (Corning 61-277-RF)) (Torr et al., 2015; Bernau et al., 2017). Alveolar lung tissue was minced and plated onto 10-cm plates (Falcon 353003) in HLF growth medium containing DMEM supplemented with 10% fetal bovine serum (FBS; HyClone, Cytiva SH30396.03), 2 mM L-glutamine (Sigma-Aldrich G7513), and antibiotics as above. Expanded populations of fibroblasts were subsequently subcultured after 4–5 days, resulting in the development of a homogenous fibroblast population. All primary cultures were used from passages 5–10.

Eosinophil isolation was performed under a study protocol approved by the University of Wisconsin-Madison Health Sciences Institutional Review Board (IRB #2013-1570). Informed written consent was obtained from subjects prior to participation. Peripheral blood eosinophils were obtained from subjects with diagnoses of allergic rhinitis and asthma (n = 3). Subjects using low doses of inhaled corticosteroids did not use their corticosteroids on the day of the blood draw.

Eosinophils were purified by negative selection as previously described (Esnault et al., 2017a, 2017b) from three different donors. Briefly, 200 ml of heparinized blood was diluted 1:1 in Hank’s Balanced Salt Solutions (HBSS; Corning 21-022-CM) and was overlaid above Percoll (1.090 g/ml; GE). After centrifugation at 700 xg for 20 min, at room temperature, the mononuclear cells were removed from the plasma/Percoll interface and erythrocytes were eliminated from the cell pellet by hypotonic lysis. The remaining pellet was resuspended in 2% new calf serum (Sigma-Aldrich; N4762) in HBSS. Cells were then incubated with anti-CD16, anti-CD3, anti-CD14 and anti-Glycophorin-A beads from Miltenyi (San Diego, CA), and run through an AutoMACS (Miltenyi). Eosinophil purity was determined by a manual count of a cytospin slide with eosin stain under the microscope. Eosinophil preparations with purity ≥ 99% were shipped on the same day, ∼5 h after the blood draw. Fresh cells were shipped overnight on ice in eosinophil culture medium containing RPMI 1640 (Corning 10041CV) supplemented with 10% FBS, 2 mM L-glutamine, and antibiotics as above using a Petaka®G3 FLAT cell culture device (Celartia, OH).

### 2.3 Cell culture

On day 1, HLFs were seeded on the inner chambers (surface area: 13.8 mm^2^, volume: 16 μL) of the coculture device at a density of 200 cells/mm^2^ (a total of 2760 fibroblasts per inner chamber) in HLF growth medium and allowed to attach and spread overnight (**Supplementary Figure 2**). On day 2, HLF media was replaced with starvation medium containing DMEM supplemented with 0.1% bovine serum albumin (BSA; Fisher BioReagents BP9706100), 2 mM L-glutamine, and antibiotics as above, and allowed to serum starve overnight. On the same day, peripheral blood eosinophils were received, retrieved, and activated in eosinophil culture medium with 2 ng/mL interleukin-3 (IL-3; BD Pharmingen 554604) at 1 × 10^6^ cells/mL for 24 h in a well plate (Corning). Human immunoglobulin G (IgG; Sigma-Aldrich I2511) was heat aggregated (HA-IgG) in phosphate buffered saline (PBS; Gibco 10010049) for 30 min at 63 °C (Esnault et al., 2017a, 2020) and used to coat outer chambers (15 μg/mL; 36 μL/chamber) of the coculture device overnight.

On day 3, HLF media was replaced with serum-free medium containing DMEM supplemented with 2 mM L-glutamine and antibiotics as above. HA-IgG was removed and outer chambers were saturated with 0.1% gelatin (Sigma-Aldrich G1890) in PBS for 30 min at 37 °C. Activated eosinophils were washed and resuspended at 2.6 × 10^6^ cells/mL in fresh eosinophil culture medium (no cytokines), then seeded to the outer chambers (surface area: 29.3 mm^2^, volume: 36 μL) at a density of 3125 cells/mm^2^ (a total of 91563 eosinophils per outer chamber). After eosinophils were seeded and allowed to attach for 1.5 h, the inner and outer chambers were connected using additional 50 μL coculture medium (1:1 ratio of HLF serum-free medium and eosinophil culture medium) to start coculture (**Supplementary Figure 2**). The coculture was allowed for 72 h without media change. In control conditions (*i*.*e*., monoculture or without HA-IgG), all procedures were performed the same way minus the one component, as indicated in figures.

The coculture period was determined based on previous studies where eosinophil-conditioned media was added to HLFs for 72 h (Esnault et al., 2017a; Bernau et al., 2018, 2021). Moreover, 72 h in coculture allows sufficient time for the signaling factors from the degranulating eosinophils to reach the HLF compartment and for HLFs to express and release cytokines to the coculture media. The seeding density of eosinophils in the microfluidic device was also translated from previous studies by keeping the surface area seeding density consistent (**Supplementary Table 1**) (Esnault et al., 2017b). Fibroblast seeding densities (157 cells/mm^2^) utilized in previous publications (Bernau et al., 2018, 2021) were modestly expanded (200 cells/mm^2^) to increase the output for assays in our microfluidic coculture devices and was examined by visual inspection after 72 h in culture.

Throughout culturing, the devices were kept in a primary container (OmniTray, Thermo Scientific 264728), where sacrificial water droplets (about 1 mL) were pipetted around the device chip to mitigate evaporation. The primary container was placed in a secondary container (BioAssay dishes, Thermo Scientific 240835), in which the peripheral of the primary container was wrapped tightly with Kimwipes (Kimberly-Clark Professional 34256) soaked with water (about 100 mL) (**Supplementary Figure 3**). The secondary container was then placed on the bottom shelf of an incubator at 37 °C and 5% CO_2_. As an additional precaution, we also alternated condition layouts for each presented experiment to mitigate potential edge effect.

### 2.4 Microscopy

For live-cell imaging, the coculture devices were placed into an OmniTray with a 50 × 61 mm rectangle cut out of the bottom and resealed with 48 × 60 mm No. 1 glass coverslips (Gold Seal® Cover Glasses, thickness 0.13-0.17 mm, Thermo Scientific 48×60-1-002G). Phase-contrast images of eosinophils were acquired using a ZEISS Primovert inverted microscope (40× objective, LD Plan-ACHROMAT 40×/0,50 Ph1 ∞/1,0; Carl Zeiss) mounted with an MU1403B Microscope Digital Camera (AmScope).

### 2.5 Enzyme-linked immunosorbent assay

After chambers were connected for 24 h, 7 μL of conditioned media from each device were harvested. Cells were removed by centrifugation at 1,500 rpm (about 1,000 g) for 10 min at 4 °C, and 5 μL of each supernatant were collected and stored at -80 °C until use. The human EDN enzyme-linked immunosorbent assay (ELISA) kit (MBL International 7630, Woburn, MA) has a minimum detection limit of 0.62 ng/mL. Samples were diluted 1:30∼1:50 with the Assay diluent and centrifuged again before use according to the manufacturer’s protocol. The absorbance was read at 450 nm (reference at 620 nm) using a Cytation 5 Cell Imaging Multi-Mode Reader (Agilent Biotek).

### 2.6 Reverse transcription quantitative PCR

HLFs were lysed at the end of experiments (72 h in connection) and the cell lysates were stored in RNase-free buffer at -80 °C until use. HLF mRNA was extracted from lysates using the Dynabeads™ mRNA DIRECT™ Micro Purification Kit (Invitrogen™ 61021). The reverse transcription reaction was performed using the High-Capacity RNA-to-cDNA™ Kit (Applied Biosystems™ 4387406). Gene expression levels were determined by qPCR using SsoAdvanced SYBR Green Supermix (Bio-Rad 1725271) in a CFX Connect Real-Time PCR Detection System (Bio-Rad Laboratories, Hercules, CA). The primer pair sequences are shown in **Supplementary Table 2** with glucuronidase β (*GUSB*) serving as a housekeeping gene (Esnault et al., 2017a).

Standard curves and primer efficiencies were previously determined (Bernau et al., 2021). Data are expressed as fold changes of control using the comparative cycle threshold (ΔΔCt) method where ΔCt = Ct of the target gene (*e*.*g*., *IL6*) – Ct of the housekeeping gene (*GUSB*); ΔΔCt = ΔCt of treated samples (*e*.*g*., “HLF + EOS (IL-3 IgG)”) – ΔCt of control samples (“HLF (IL-3 IgG)”); and fold change = 2^-ΔΔCt^ (Esnault et al., 2017b).

### 2.7 Multiplex immunoassays

At the end of experiments, all culture supernatants were harvested from each device (about 95 μL). Cells were removed by centrifugation at 1,500 rpm (about 1,000 g) for 10 min at 4 °C, and 75 μL of each supernatant were collected and stored at -80 °C until use. Soluble factor analyses were performed using a custom multiplex immunoassay panel for IL-6, IL-8/CXCL8, and GM-CSF (MILLIPLEX HCYTA-60K, MilliporeSigma). Samples were centrifuged again before use according to the manufacturer’s protocol. Assays were read on a Luminex 200 instrument (Luminex Corp.) with xPONENT software. Standard curves were obtained using Belysa® Immunoassay Curve Fitting Software (MilliporeSigma). Data below the limit of quantification were plotted as zero on graphs.

### 2.8 Statistical analyses

Data are expressed as the mean of culture (technical) replicates ± the standard error of the mean (SEM) (n=3-4); each plotted point represents a culture replicate within the same device chip. Data were analyzed and visualized using Graphpad Prism 9 software (San Diego, CA). Differences among culture conditions were analyzed using ordinary one-way analysis of variance tests (one-way ANOVA), followed by Holm-Šídák’s multiple comparisons test to compare the means of preselected pairs of conditions. p < 0.05 was considered statistically significant.

## 3. Results and Discussion

### 3.1 Design and workflow of the open microfluidic coculture device

In this study, we present an open microfluidic coculture model to investigate the bidirectional signaling between HLFs and blood eosinophils and their roles in airway inflammation. To establish this *in vitro* model, we adapted a previously developed open microfluidic coculture device made of polystyrene, where two culture chambers are separated by a half wall but can be later connected to start coculture (Zhang et al., 2020). Our device is purposefully designed to be simple and easy to use, which are factors that are increasingly desired in microfluidic devices to enable adoption by biology labs (in contrast to devices with complicated pumps and peripheral equipment) (Sackmann et al., 2014). The coculture devices are fabricated in an array to allow for testing of multiple experimental conditions in one chip, and a chip is designed to fit in a standard 10-cm petri dish, which can be used as a primary container for culture sterility and evaporation control (**Figure 1A**). The petri dish (and a secondary container) can then be fitted easily into an incubator and imaged with phase contrast or fluorescence microscopy directly using existing microscope holders.

Each of the microfluidic coculture devices is composed of an inner and an outer chamber, separated by a half wall (**Figure 1B**). The chambers are designed to be operated simply with a standard pipette; the pipette tip (up to the size of a P200) can be anchored to the notch, and the liquid will fill the entire culture chamber with a single dispensing step due to the spontaneous capillary flow (Casavant et al., 2013; Berthier et al., 2019; Zhang et al., 2020). Also, because the chambers are on the same focus plane (unlike Transwell^®^ inserts where the membrane is higher than the well plate bottom), the two cell types in coculture can be easily imaged at the same time using a standard microscope. The compatibility of our coculture devices with traditional workflows (*e*.*g*., use of petri dishes, pipettes, and microscopes) allows them to be used conveniently by most biologists.

To demonstrate the use of coculture devices in the study, a simplified workflow of one of the experimental conditions (“HLF + EOS (IL-3 IgG)”) is shown (**Figure 1C, Supplementary Figure 2**). Briefly, HLFs were first seeded to the inner chamber of the coculture device, allowing them to attach and spread out. Next, eosinophils activated by interleukin-3 (IL-3) were seeded to the outer chambers which were previously coated with heat-aggregated immunoglobulin (HA-IgG). After the eosinophils attached, the inner and outer chambers were connected with additional coculture medium to overflow the half wall and initiate coculture. The two cell types were allowed to communicate via soluble factors for 72 h.

### 3.2 Eosinophils degranulate on HA-IgG in the microfluidic coculture device

Data are expressed as mean ± SEM (n=4); each point is one culture replicate. Data were analyzed using one-way ANOVA, followed by Holm-Šídák’s multiple comparisons test.

Eosinophil degranulation starts with the release of soluble factor contents, such as toxic proteins (*e*.*g*., EDN), from intracellular granules while eosinophils are alive (Spencer et al., 2014).

Adhesion of activated eosinophils typically triggers degranulation (Cook et al., 2004; Lintomen et al., 2008). In contrast, following eosinophil cell death (*e*.*g*., apoptosis or cytolysis), cell-free granules and other cellular contents are released as the cell membrane ruptures (Esnault et al., 2020). Additionally, it has been shown that extracellular cell-free granules also contain surface receptors that allow them to be activated and to further degranulate even when outside of cells (Neves et al., 2008); thus, cell-free granules from eosinophils can also respond to soluble factors secreted by neighboring cells (*e*.*g*., fibroblasts).

We previously developed an *ex vivo* eosinophil degranulation model that recapitulates the responses of activated airway eosinophils to a biologically relevant extracellular cue found in the human lung airways (Esnault et al., 2017b). Briefly, freshly isolated blood eosinophils were primed with IL-3 and seeded to well plates coated with or without HA-IgG. Eosinophil degranulation occurred as soon as 30 min after adhesion to HA-IgG, and significant cytolysis was observed after 4 h (Esnault et al., 2020). To translate the eosinophil degranulation model established on well plates to the microfluidic coculture device, the first step is to validate that primary eosinophils will behave similarly in the coculture devices.

In accordance with our previous work (Esnault et al., 2020), after being seeded to the outer chambers for 1.5 h, eosinophils pre-activated by IL-3 displayed apparent adhesion and spreading on HA-IgG, while still maintaining their cellular membrane integrity (red arrows, **Figure 2A**: “EOS (IL-3 IgG)” or “EOS (IL-3 IgG) + HLF”). In contrast, eosinophils seeded to devices without HA-IgG appeared in suspension and some displayed polarized morphology due to the prior activation with IL-3 (**Figure 2A**: “EOS (IL-3)” or “EOS (IL-3) + HLF”) (Esnault et al., 2017b; Shen et al., 2021). After the chambers were connected for 24 h and as expected according to our previous work (Esnault et al., 2020), IL-3 IgG-activated eosinophils had likely undergone significant cytolysis, leaving aggregates of cell debris (yellow asterisks, **Figure 2A**). At 72 h, control eosinophils not exposed to HA-IgG also released some cell-free granules (white arrowheads, Figure 2A), suggesting eosinophil death after prolonged IL-3-free culture *ex vivo* (Vancheri et al., 1989; Esnault et al., 2015). Images from eosinophil donor 1/HLF donor 2 are shown and are representative of a total of 6 donor pairs (eosinophil donor 1-3, HLF donor 1 and 2); images from all donor pairs can be found in **Supplementary Figure 4**.

**FIGURE 2.**
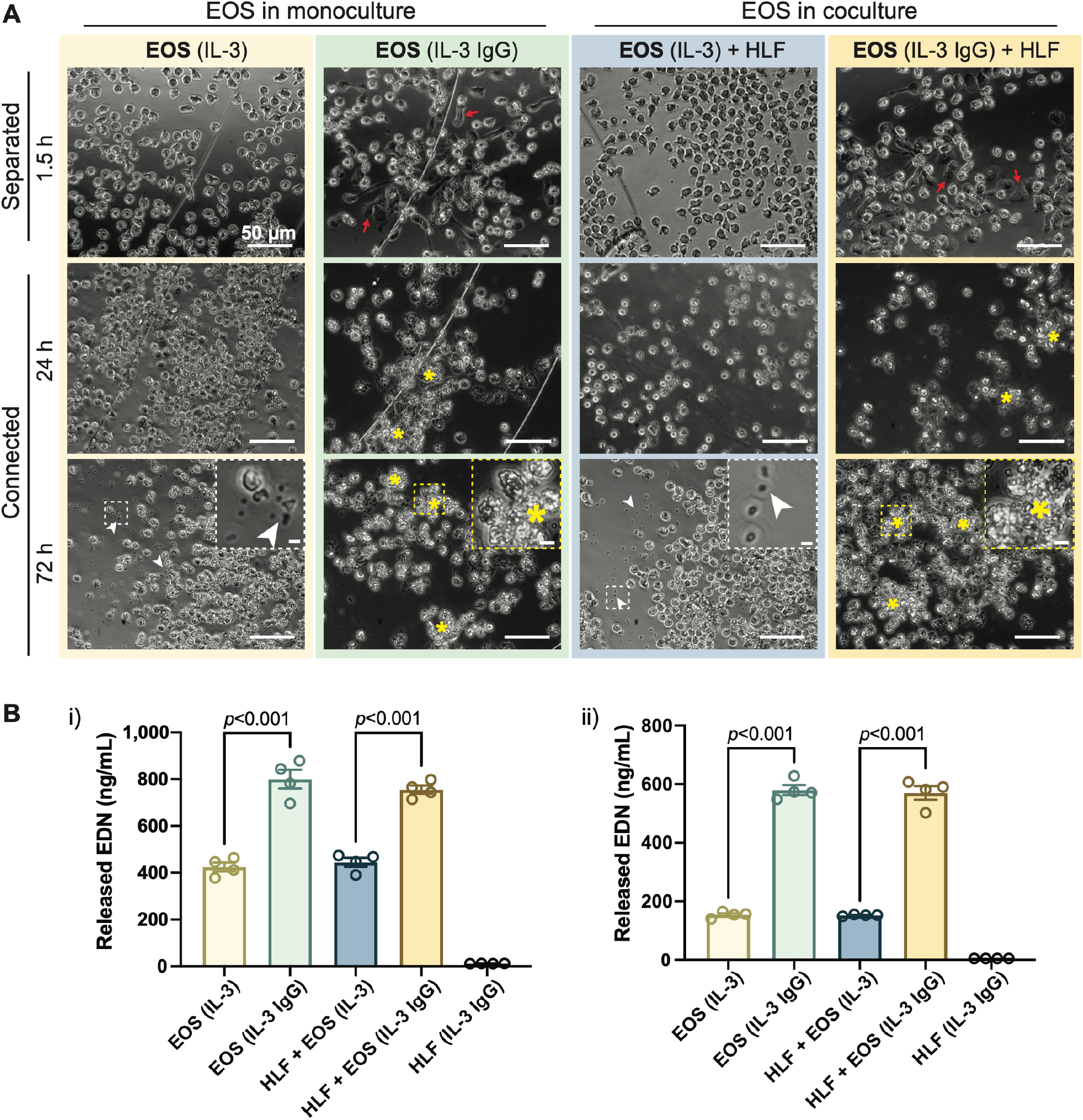
IL-3-activated eosinophils degranulate on HA-IgG in the coculture device. **(A)** Phase contrast images of eosinophils (EOS) activated with IL-3 (“IL-3”) and in some conditions also seeded onto HA-IgG (“IL-3 IgG”) in coculture devices at the indicated times. The chambers remained separated for 1.5 h and then were connected for up to 72 h. Red arrows show eosinophils spreading on HA-IgG. Yellow asterisks show aggregates of eosinophil cell debris. White arrowheads show eosinophil cell-free granules. Magnified inserts included for improved visualization of cell debris and cell-free granules (scale bars: 50 μm). Images are representative of four culture replicates, as well as three eosinophil donors and two HLF donors. **(B)** Levels of released eosinophil-derived neurotoxin (EDN) in culture media after 24 h measured by ELISA. Results from two different representative eosinophil/HLF donor pairs are shown in **i**) and **ii**).

Because both chambers of the open microfluidic coculture device are on the same focus plane, we were able to monitor HLFs together with eosinophils in culture at the same time. Across different culture conditions, we did not see obvious morphological dissimilarities of HLFs (**Supplementary Figure 5**); the HLFs maintained a spindle-like morphology (characteristic of fibroblasts) in both monoculture and coculture conditions. Further, we observed a consistent increase in cell density as the experiment progressed, indicating growth and expansion of HLFs in our microfluidic device.

Conditioned media from the connected chambers was collected after 24 h, and the EDN content measured using ELISA. Eosinophils in IL-3 IgG conditions (with or without HLF) released significantly more EDN than control eosinophils in IL-3 conditions (**Figure 2B**). Data from eosinophil donor 1/HLF donor 1 (**Figure 2Bi**) and eosinophil donor 2/HLF donor 2 (**Figure 2Bii**) were shown and are representative of a total of 4 donor pairs (eosinophil donor 1 and 3, HLF donor 1 and 2); data from all available donor pairs can be found in **Supplementary Figure6.**We note that there is some donor-to-donor variation; importantly, the trends hold across all donor pairs and are statistically significant.

Taken together, our results show that IL-3-activated eosinophils further degranulate on HA-IgG-coated outer chambers, suggesting we were able to recapitulate the previously developed eosinophil degranulation model (Esnault et al., 2017b) in the open microfluidic coculture device.

### 3.3 *CXCL8, IL6*, and *ICAM1* are upregulated in HLFs cocultured with degranulating eosinophils

After the validation of the eosinophil degranulation model in the microfluidic coculture device, we wished to investigate the response of HLFs to degranulating eosinophils in coculture. First, we aimed to determine the changes of HLFs at the mRNA expression level. To achieve this goal, HLFs from the inner chambers of the coculture device were lysed after 72 h and analyzed using RT-qPCR. Notably, in HLF monoculture conditions (“HLF (IL-3 IgG)”), the coculture devices were treated the same way as in “HLF + EOS (IL-3 IgG)” conditions minus the presence of eosinophils (*i*.*e*., outer chambers were coated with HA-IgG and loaded with IL-3 treated medium before connection) to account for the possible impact of residual IL-3 and released HA-IgG from the process on HLFs.

Our previous RNA-sequencing analyses identified several genes that were upregulated in HLFs cultured with conditioned media from degranulated eosinophils, which had downstream functions on inflammation, tissue remodeling, and lipid synthesis (Esnault et al., 2017a). Among those genes, we selected 3 representative genes that promote granulocytic inflammation to measure in our study: interleukin 6 (*IL6*), C-X-C motif chemokine ligand 8 (*CXCL8*), and intercellular adhesion molecule 1 (*ICAM1*). As a result, HLFs in coculture with degranulating eosinophils (“HLF + EOS (IL-3 IgG)”) showed the highest expression of *IL6* (**Figure 3A**), *CXCL8* (**Figure 3B**), and *ICAM1* (**Figure 3C**), significantly higher than control HLFs, those in monoculture (“HLF (IL-3 IgG)”) or in coculture with non-degranulating eosinophils (“HLF + EOS (IL-3)”). These trends were consistent with our previous RNA expression results using conditioned media (Esnault et al., 2017a; Bernau et al., 2018). Data from eosinophil donor 1/HLF donor 1 **i**) and eosinophil donor 2/HLF donor 2 **ii**) were shown and are representative of a total of 4 donor pairs (eosinophil donor 1 and 2, HLF donor 1 and 2); data from all available donor pairs can be found in **Supplementary Figure 7**.

**FIGURE 3.**
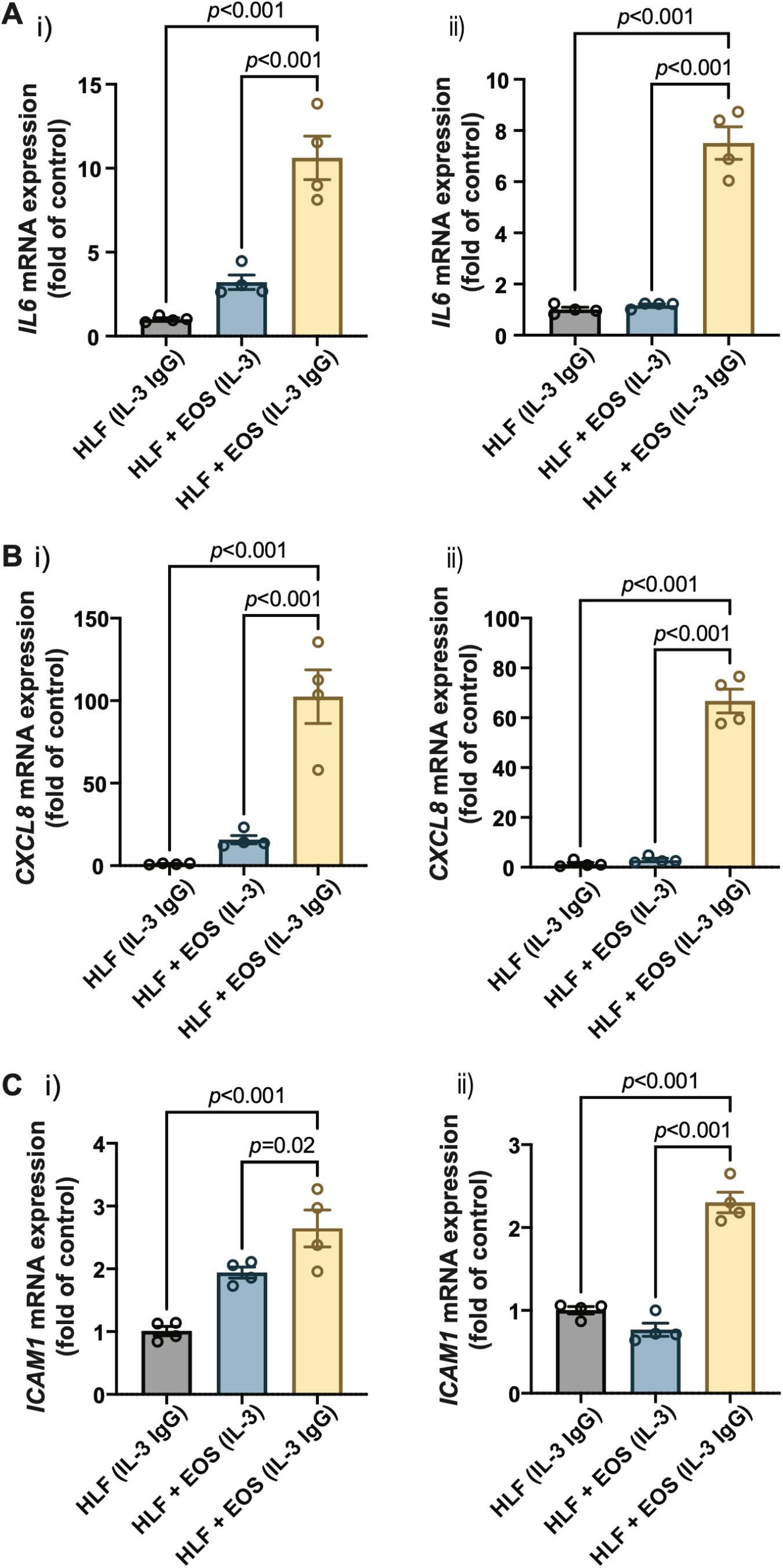
Proinflammatory genes are upregulated in HLFs cocultured with degranulating eosinophils. HLFs in coculture with degranulating eosinophils (“HLF + EOS (IL-3 IgG)”) for 72 h showed the highest mRNA level upregulation of **(A)** *IL6*, **(B)** *CXCL8*, and **(C)** *ICAM1*. Results from two representative eosinophil/HLF donor pairs are shown in i) and ii). Data are expressed as mean ± SEM (n=4); each point is one culture replicate. Data were analyzed using one-way ANOVA, followed by Holm-Šídák’s multiple comparisons test.

Together, our results demonstrate that HLFs were activated by the coculture with degranulating eosinophils in the microfluidic coculture device, as indicated by the upregulation of proinflammatory genes, *IL6, CXCL8*, and *ICAM1*.

### 3.4 Coculture of HLFs with degranulating eosinophils leads to increased secretion of IL-6 and IL-8 than in either HLF or EOS monoculture

Next, we sought to assess the cellular changes at the protein expression level, specifically the level of secreted (extracellular) proteins. After the eosinophils and HLFs were cultured in the microfluidic coculture device for 72 h, we collected the culture supernatants and analyzed their contents of IL-6 and IL-8, proteins encoded by the two genes characterized above: *IL6* and *CXCL8*, respectively. IL-6 and IL-8 are important proinflammatory cytokines as they play vital roles in neutrophil regulation, with IL-8 as a neutrophil chemoattractant (Hammond et al., 1995) and IL-6 as a neutrophil activator (Fielding et al., 2008).

Similar to the results reported previously (Bernau et al., 2018, 2021), we observed a significant increase of both IL-6 (**Figure 4A**) and IL-8 (**Figure 4B**) in the coculture conditions of HLFs with degranulating eosinophils (“HLF + EOS (IL-3 IgG)”) compared to the control conditions (*i*.*e*., monoculture or non-degranulating eosinophils). Notably, the trends of IL-6 and IL-8 release among the three conditions with HLFs were similar to the trends of *IL6* (**Figure 3A**) and *CXCL8* (**Figure 3B**) mRNA expression, suggesting the changes of the two markers took place at both transcription and translation levels in HLFs. Data from eosinophil donor 1/HLF donor 1 **i**) and eosinophil donor 3/HLF donor 2 **ii**) were shown and are representative of a total of 6 donor pairs (eosinophil donor 1-3, HLF donor 1 and 2); data from all donor pairs can be found in **Supplementary Figure 8**.

**FIGURE 4.**
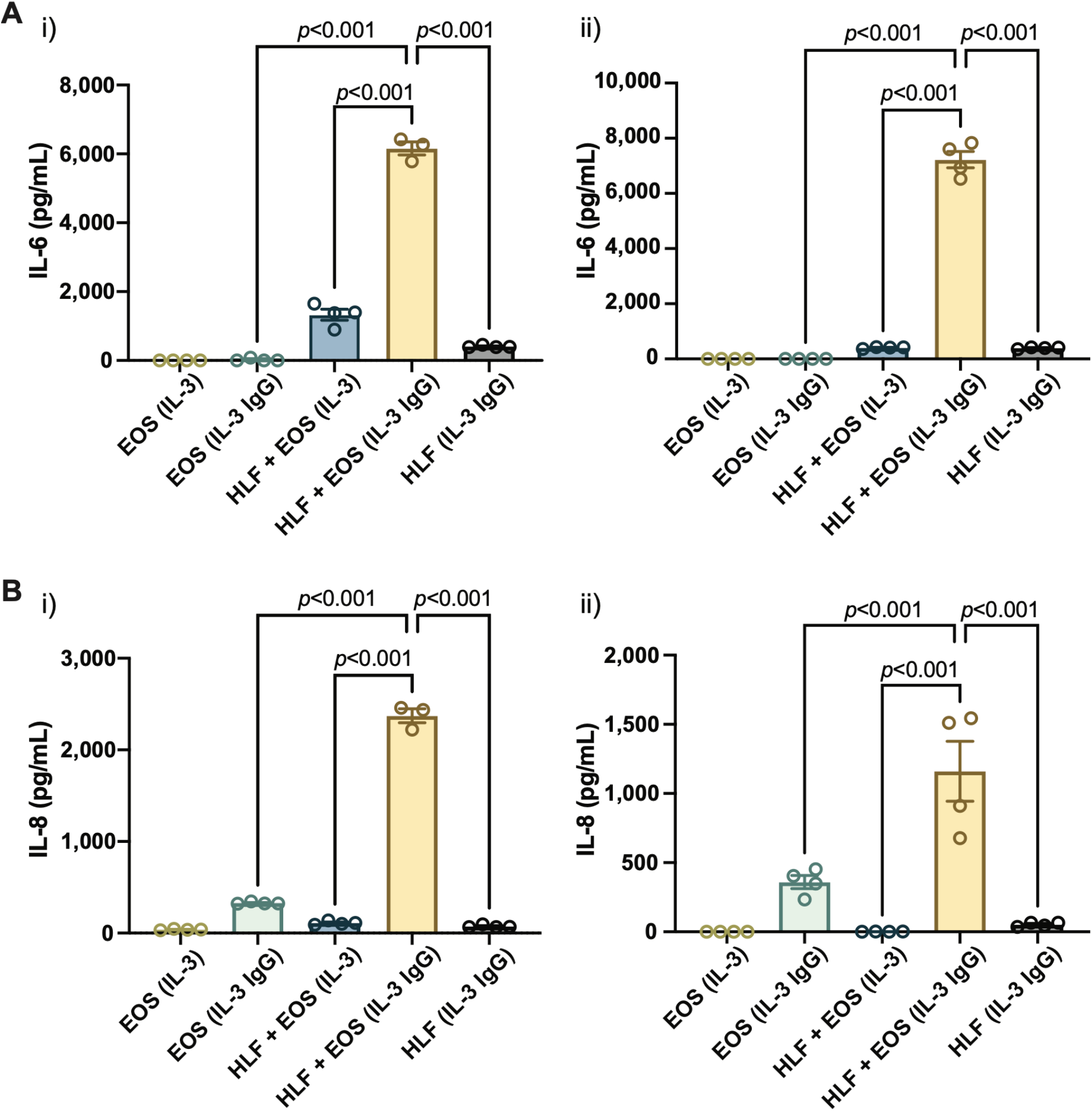
The coculture of HLFs with degranulating eosinophils releases the highest levels of proinflammatory cytokines compared to HLF and eosinophil monocultures and other control conditions. Levels of soluble factors, **(A)** IL-6 and **(B)** IL-8, were measured in the conditioned media after 72 h in culture. Results from two representative eosinophil/HLF donor pairs are shown in **i**) and **ii**). Data are expressed as mean ± SEM (n=3-4); each point is one culture replicate. Data were analyzed using one-way ANOVA, followed by Holm-Šídák’s multiple comparisons test.

Together, our results suggest that the coculture of HLFs with degranulating eosinophils induced a proinflammatory microenvironment in the microfluidic coculture device, as indicated by the increased levels of IL-6 and IL-8 compared to either cells in monoculture or the coculture with non-degranulating eosinophils.

### 3.5 GM-CSF expression and secretion are increased in the coculture of HLFs with degranulating eosinophils

Having validated our coculture model with previously reported analytes from degranulating eosinophils that activate HLFs, we then sought to discover additional soluble factors that could have more bidirectional effects on the eosinophil-HLF crosstalk. GM-CSF has been found to enhance eosinophil survival *in vitro* and accumulation *in vivo*, even at a minute level (Esnault and Malter, 2001; Nobs et al., 2019). Research shows that fibroblasts can promote eosinophil survival and adherence via the increased secretion of GM-CSF, but the studies were done in direct coculture systems (with cell-cell contact) (Vancheri et al., 1989; Zhang et al., 1996; Solomon et al., 2000). Further, the primary cell sources used in the previous studies were different from ours: we used normal HLFs and blood eosinophils from donors with allergy and asthma; Vancheri (1989) used normal HLFs and blood eosinophils from donors with allergic rhinitis; Zhang (1996) used bronchial myofibroblasts and blood eosinophils from donors with high eosinophil counts; Solomon (2000) used conjunctival fibroblasts and blood eosinophils from donors with mild atopy. Herein, we aimed to study if GM-CSF could be upregulated in our segregated coculture system (without cell-cell contact) with our HLFs and eosinophils.

Similar to other analytes presented in the study, we first characterized the mRNA expression of colony stimulating factor 2 (*CSF2*), the protein encoding gene of GM-CSF, in HLFs that were in coculture or monoculture for 72 h in the microfluidic device. According to the results (**Figure 5A**), the expression of *CSF2* was upregulated the most in HLFs cocultured with degranulating eosinophils (“HLF + EOS (IL-3 IgG)”). Then, we measured the levels of soluble GM-CSF secreted into the culture supernatants after 72 h and observed a similar trend (**Figure 5B**): the release of GM-CSF is significantly higher in the coculture conditions of HLFs with degranulating eosinophils than in monocultures or cocultures with non-degranulating eosinophils. Data from eosinophil donor 2/HLF donor 1 **i**) and eosinophil donor 2/HLF donor 2 **ii**) were shown and are representative of up to 6 donor pairs; data from all available donor pairs can be found in **Supplementary Figure 9**.

**FIGURE 5.**
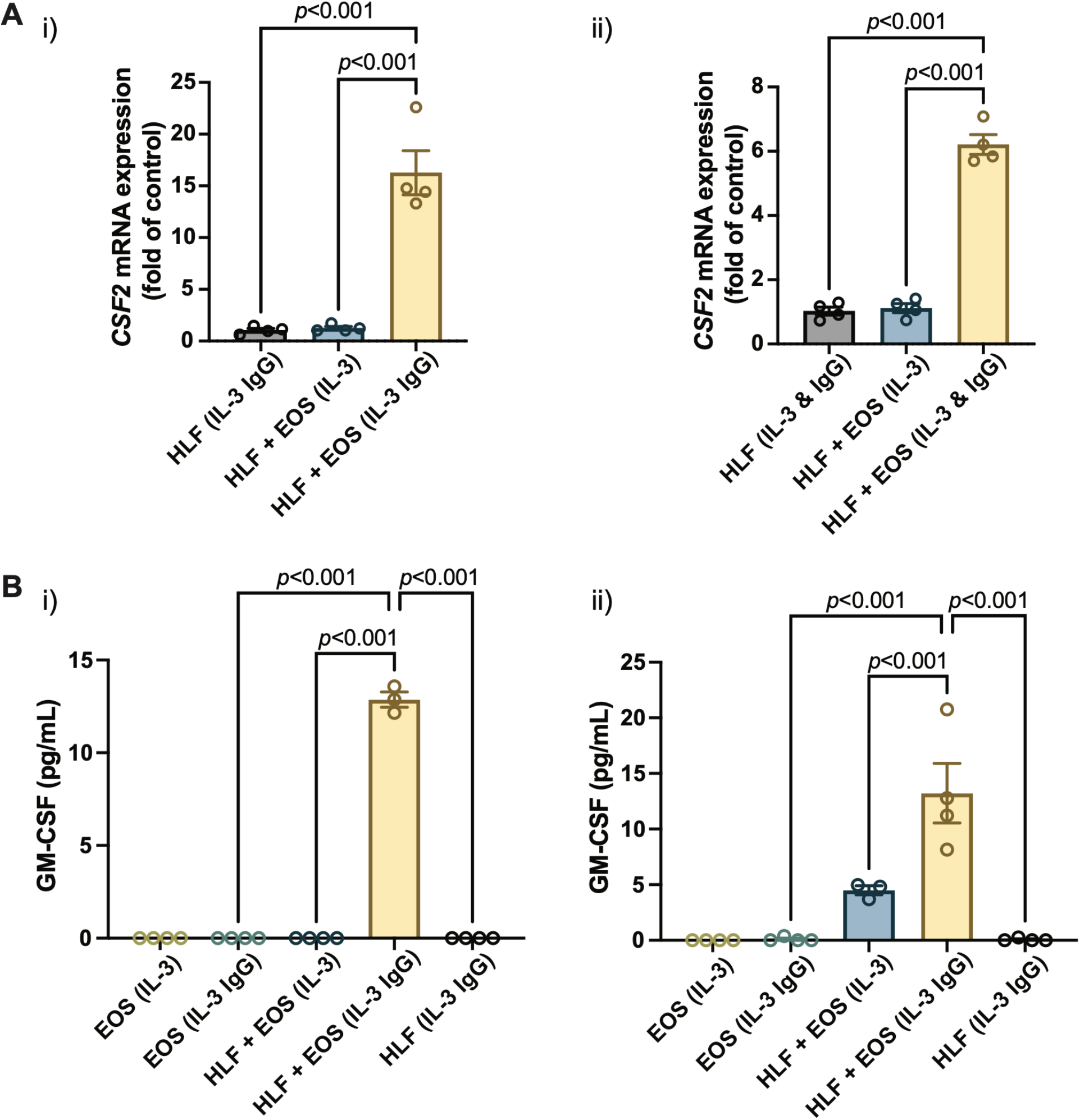
The coculture of HLFs with degranulating eosinophils induces the highest levels of **(A)** mRNA expression of CSF2 in HLFs and **(B)** protein secretion of GM-CSF in the conditioned media after 72 h in culture. Results from two representative eosinophil/HLF donor pairs are shown in **i**) and **ii**). Data are expressed as mean ± SEM (n=3-4); each point is one culture replicate. Data were analyzed using one-way ANOVA, followed by Holm-Šídák’s multiple comparisons test.

Jointly, the results suggest that the GM-CSF expression was upregulated in HLFs when in coculture with degranulating eosinophils. The increased levels of GM-CSF in coculture media could in turn prolong the survival of eosinophils in our *ex vivo* coculture system as a bidirectional signaling effect from HLFs. This new finding, to our knowledge, is the first to show GM-CSF upregulation in a segregated coculture system of eosinophils and fibroblasts; it could provide us with further directions in studying the impacts from HLFs to eosinophils, given the importance of GM-CSF in eosinophil activation *in vitro* and *in vivo*.

Despite many benefits (enabling bidirectional signaling, *etc*), we acknowledge the limitation of the shared media mechanism of our microfluidic coculture devices: as with other coculture systems, the shared media can lead to ambiguity when determining the source of the soluble factors in coculture. Consequently, complementary studies that characterize only one type of cells (*e*.*g*., conditioned media, qPCR, microscopy) are needed to fully investigate the signaling mechanisms. For example, in our study, because the culture media was shared between the degranulating eosinophils and HLFs in coculture, we cannot definitively pinpoint the source of the elevated levels of soluble factors (*i*.*e*., IL-6, IL-8, GM-CSF) in coculture, since either cells could be responsible for this effect. However, considering the findings in previous conditioned media studies (Vancheri et al., 1989; Esnault et al., 2017a; Bernau et al., 2018, 2021) and our RT-qPCR data as above (i.e., *IL6, CXCL8, CSF2*), which explicitly analyzed HLFs only, we suspect that the HLFs are likely the predominant source of the additional IL-6, IL-8, and GM-CSF in our coculture model. Still, more experiments are needed (*e*.*g*., transcriptomic studies of eosinophils) before a definitive conclusion is drawn.

Taken together, our mRNA (**Figure 3 and 5A**) and secreted protein results (**Figure 4 and 5B**), in conjunction with previous publications (Vancheri et al., 1989; Esnault et al., 2017a; Bernau et al., 2018, 2021), suggest that the HLFs in coculture with degranulating eosinophils likely secrete increased levels of IL-6, IL-8, and GM-CSF, compared to the coculture with non-degranulating eosinophils or either cell type in monoculture. This suggests the induction of HLFs into a proinflammatory phenotype in the presence of degranulating eosinophils in the microfluidic coculture device, which could in turn further activate eosinophils and amplify the inflammatory response.

## 4. Conclusion

In summary, we demonstrated the use of a novel *ex vivo* microfluidic coculture model of degranulating eosinophils and HLFs to investigate the mechanisms of airway inflammation. Our open microfluidic coculture devices present a unique platform for interrogating these cell interactions, as a method of modeling airway inflammation and fibroblast activation *ex vivo*. In this model system, IL-3-activated eosinophils strongly degranulate on HA-IgG and induce HLFs into a proinflammatory phenotype via coculture. Here, we validated the findings from our previous eosinophil-conditioned media studies; with the signaling molecules we characterized in this study (*i*.*e*., IL-6, IL-8), we found similar trends to our previous unidirectional culture system. In addition, we presented the results of a new analyte not reported before (*i*.*e*., GM-CSF), the upregulation of which could provide us with further directions in bidirectional signaling effects, especially from HLFs to eosinophils. In the future, we will continue discovering additional signaling molecules that may only be captured in our coculture (bidirectional signaling) system. Further, we plan to reengineer the device to enable triculture with additional relevant immune cells.

Our coculture model suggests an important interplay between the immune (*e*.*g*., eosinophils) and mesenchymal (*e*.*g*., HLFs) compartments leading to a proinflammatory fibroblast phenotype that may in turn amplify the overall inflammatory response. More importantly, although the present study uses eosinophils derived from people with allergy and asthma, in the future, we envision that this coculture model system could be more broadly applied to other organs (*e*.*g*., liver, breast, prostate, gastrointestinal tract) with immune-mesenchymal cell interactions in both normal and diseased states.

## 6. Author contributions

YZ, SE, KB, and ABT conceptualized the research. KB and ABT supervised the research. YZ, XS, SE, KB, and ABT planned the experiments. JWP and SJS advised on cell culture in the microfluidic coculture device. FYL, LCD, NNJ, SKM, NS, EB, SE, KB, and ABT provided expertise in the isolation and culture of eosinophils and fibroblasts, overall microfluidic device design/assay development, and the clinical significance of the work. YZ and MGT fabricated the devices. UNL optimized the solvent bonding process. PSF isolated the blood eosinophils. YZ, XS, and MGT conducted experiments and collected data. YZ analyzed the data. YZ, SE, KB, and ABT interpreted the results of experiments. YZ, MGT, UNL, KB, and ABT drafted the manuscript. YZ prepared the figures. YZ, FYL, NNJ, SKM, SE, KB, and ABT edited the manuscript. All authors reviewed and approved the manuscript.

## 8. Acknowledgements

We thank Dr. Tianzi Zhang for guidance on the operation of the coculture device. We are grateful for the collaboration with the UW-Madison Carbone Cancer Center Translational Science Biocore Biobank that led to obtaining of de-identified lung tissue specimens.

