## Supplementary for "An Open Microfluidic Coculture Model of Fibroblasts and Eosinophils to Investigate Mechanisms of Airway Inflammation"

### *Supplementary Material*

#### 1 Supplementary Figures

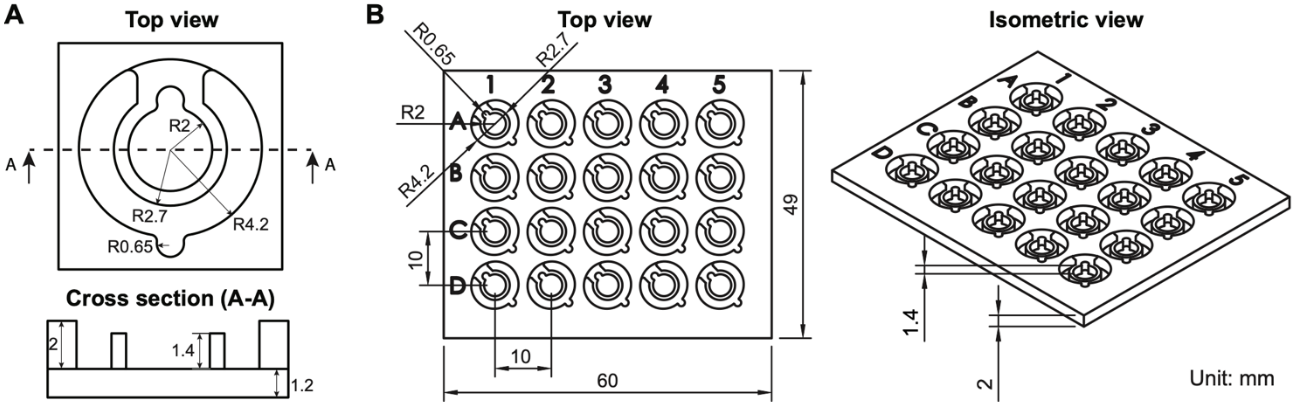


**Supplementary Figure 1.** Detailed dimensions of the open microfluidic coculture device. **(A)** Dimensions of a single coculture device. Reproduced from Zhang et al., (2020) as allowed per journal policy. Computer aided design file can be found in the Supplementary Material of Zhang et al. **(B)** Dimensions of a milled polystyrene top piece with the coculture devices in a 4×5 array.

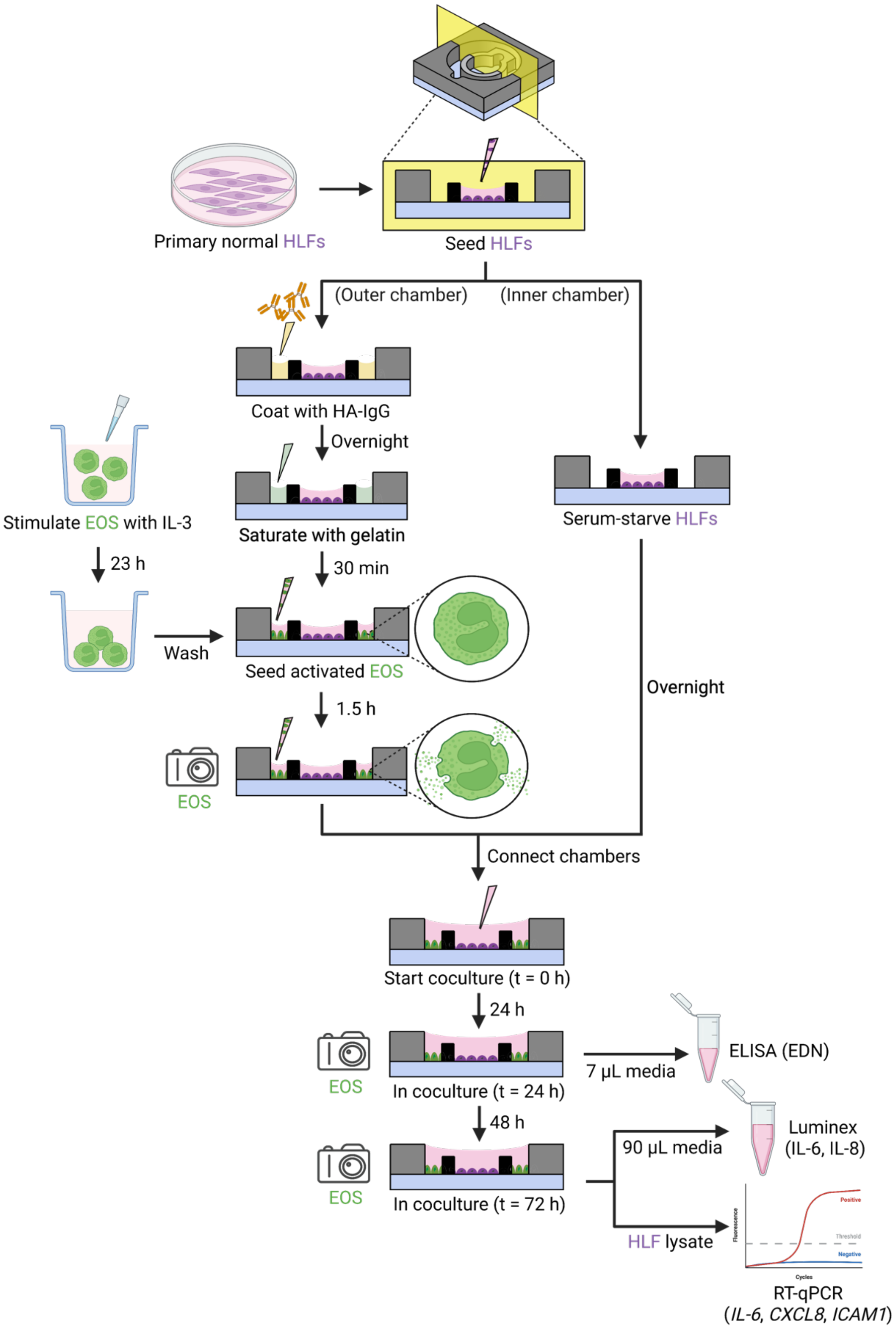


**Supplementary Figure 2.** Schematic workflow showing cell culture procedures and readouts of the eosinophil (EOS)-human lung fibroblast (HLF) coculture in the open microfluidic coculture device. Created with BioRender.com. HA-IgG: heat-aggregated immunoglobulin G. ELISA: enzyme-linked immunosorbent assay. EDN: eosinophil-derived neurotoxin. IL-6: interleukin-6. IL-8: interleukin-8.

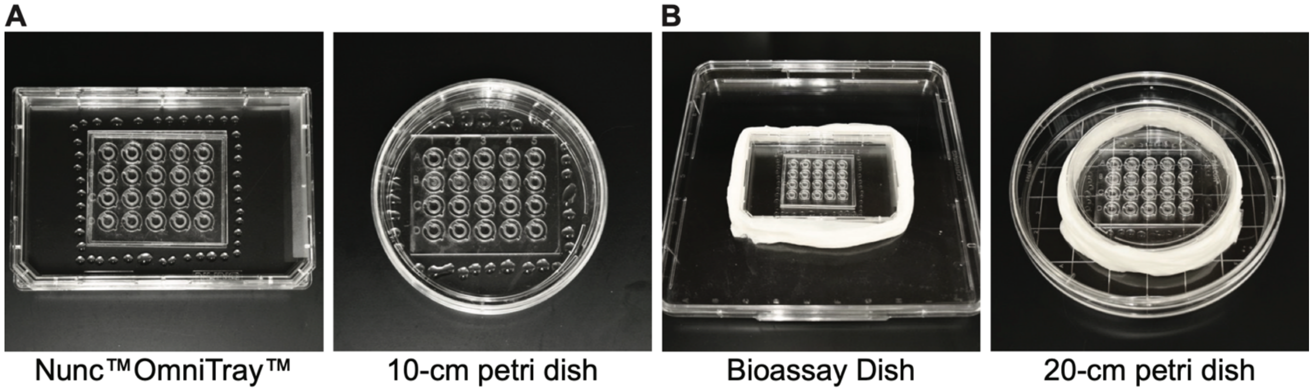


**Supplementary Figure 3.** Examples of the evaporation control setup for the open microfluidic device. **(A)** Approximately 1 mL of sterile sacrificial water was placed as droplets evenly around the perimeter of the device chip in the primary container (*e.g.,* OmniTray$^{TM}$, 10-cm petri dish). **(B)** The primary container was placed into a secondary container (*e.g.,* 30×30 cm bioassay dish or 20-cm petri dish). The perimeter of the primary container was wrapped tightly with Kimwipes soaked with sterile water. In this study, we used OmniTray$^{TM}$ and Bioassay Dish as the primary and secondary container, respectively.

*In the next pages, we show a complete collection of results from all donor pairs (including the 2 pairs shown in the main manuscript). Figure captions might be on a different page due to large figures.*

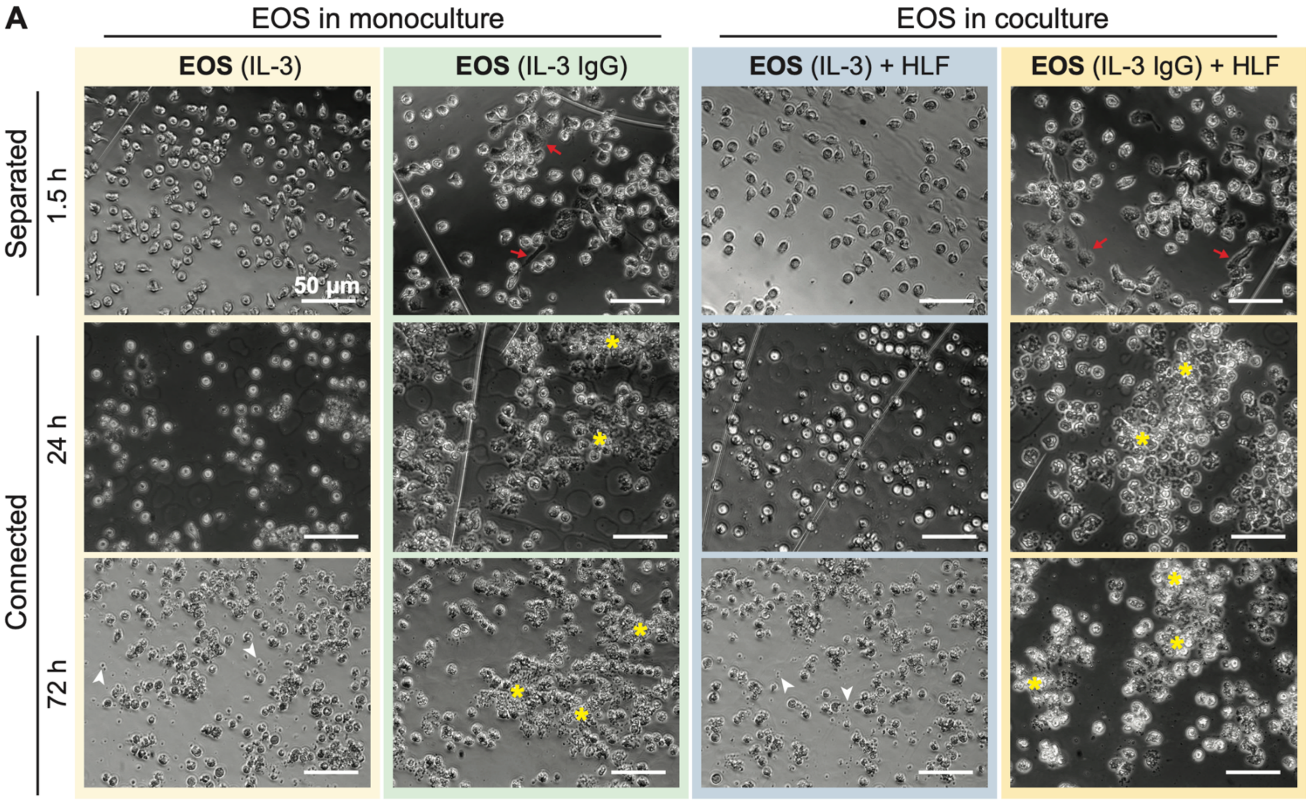


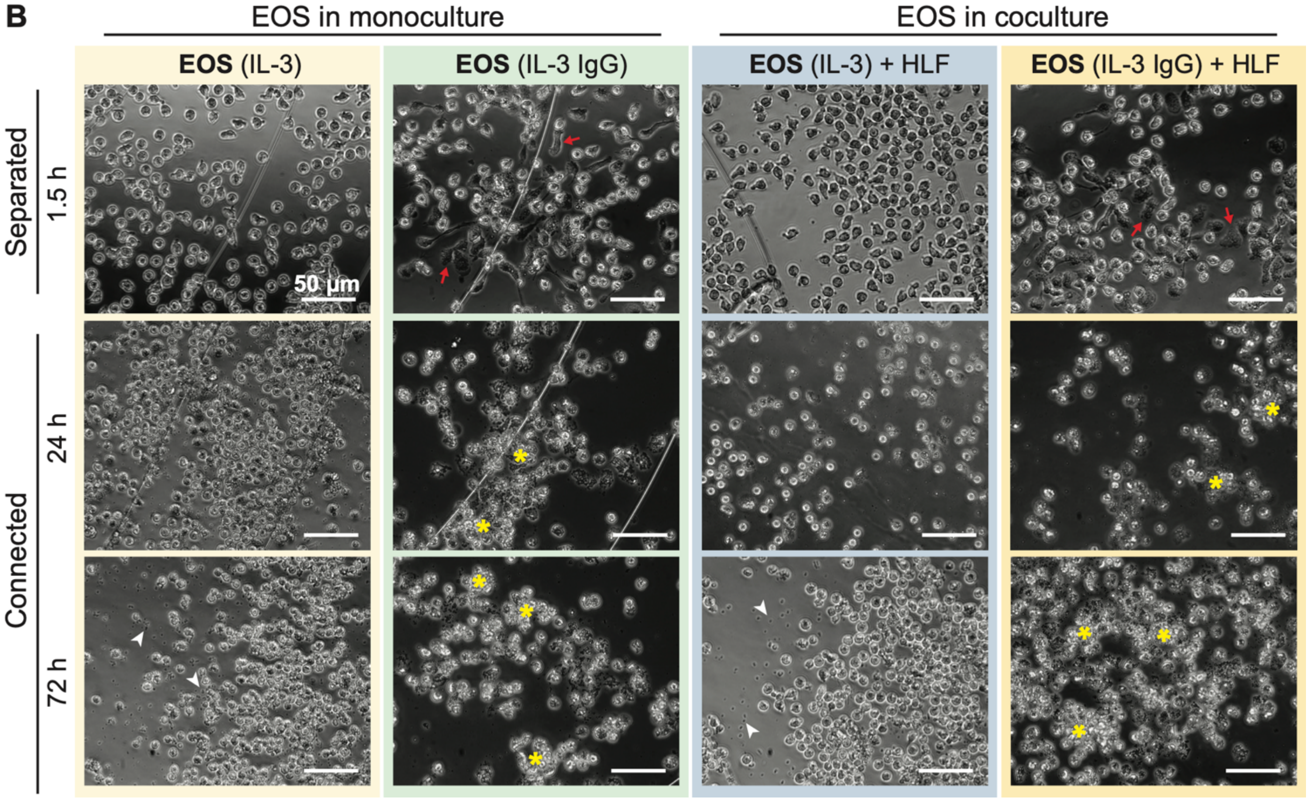


(continued next page)

C

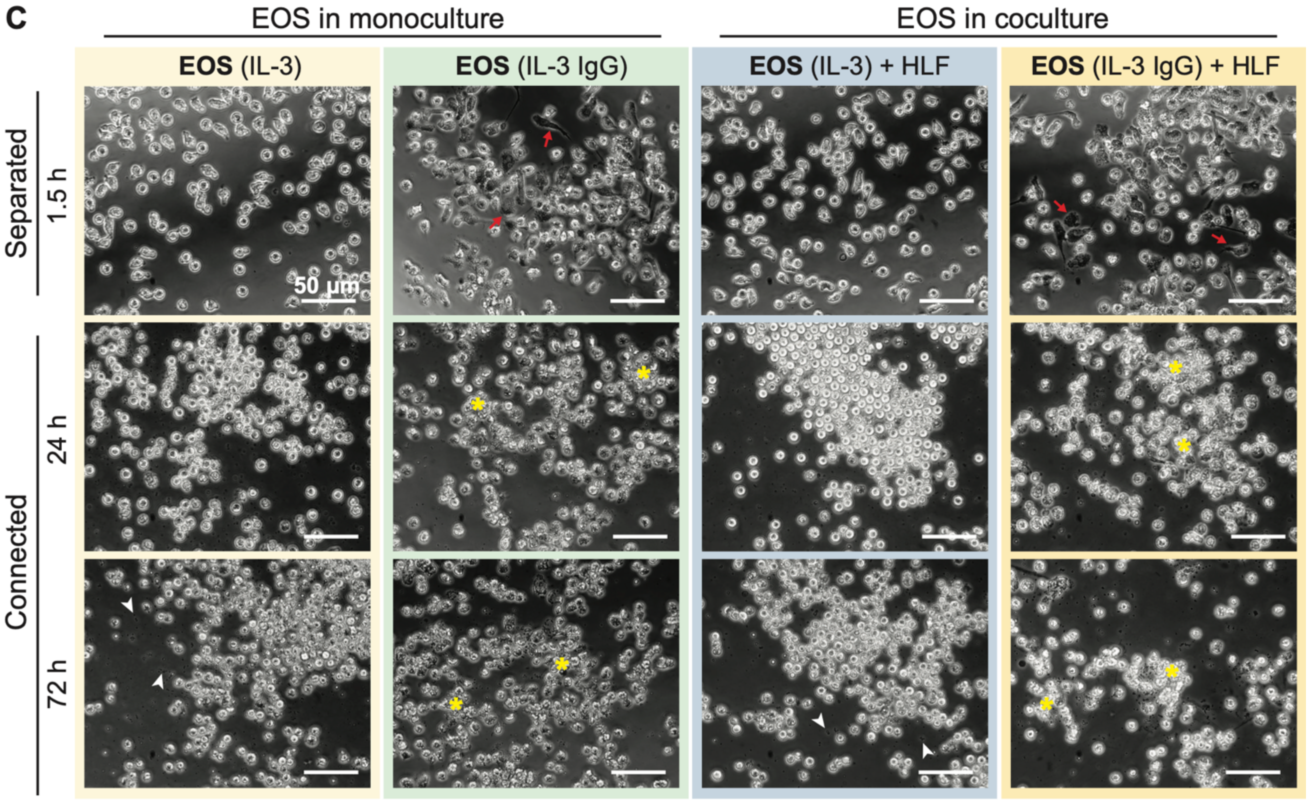


D

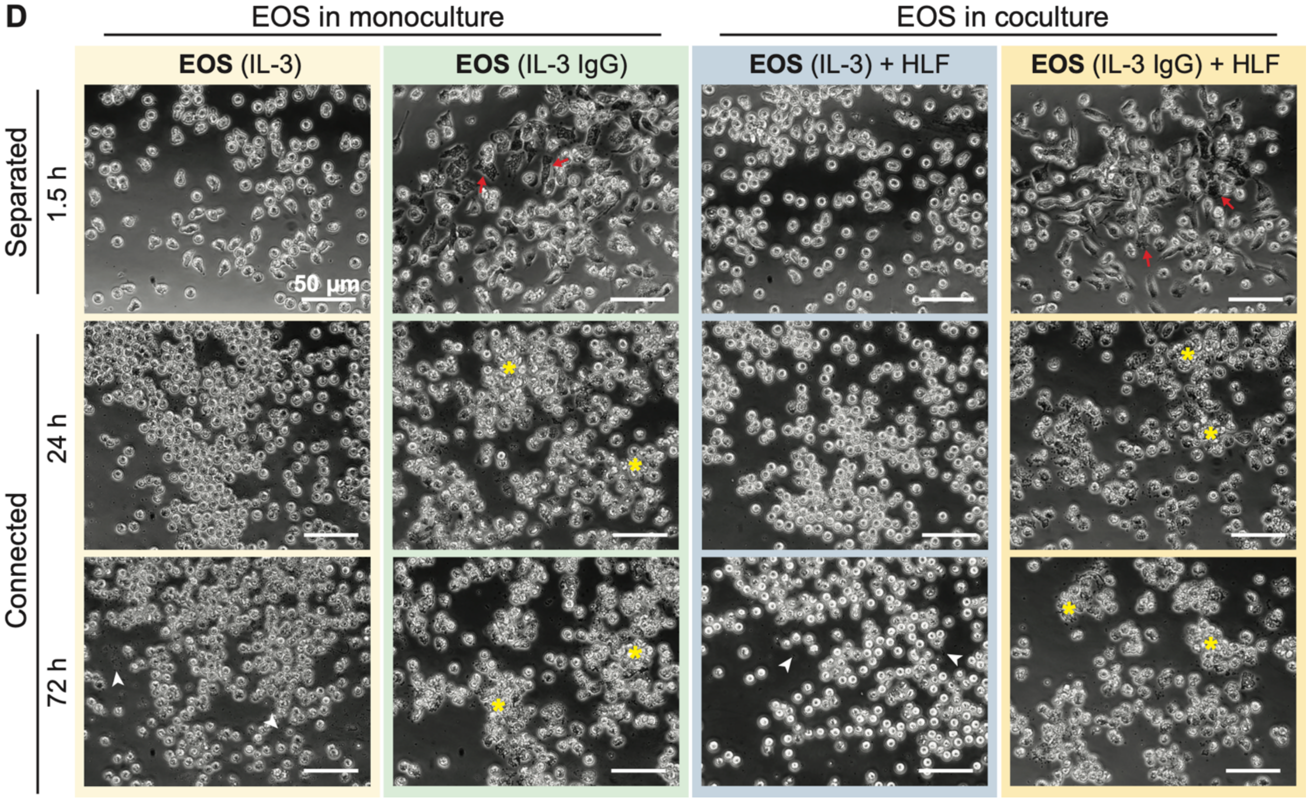


(continued next page)

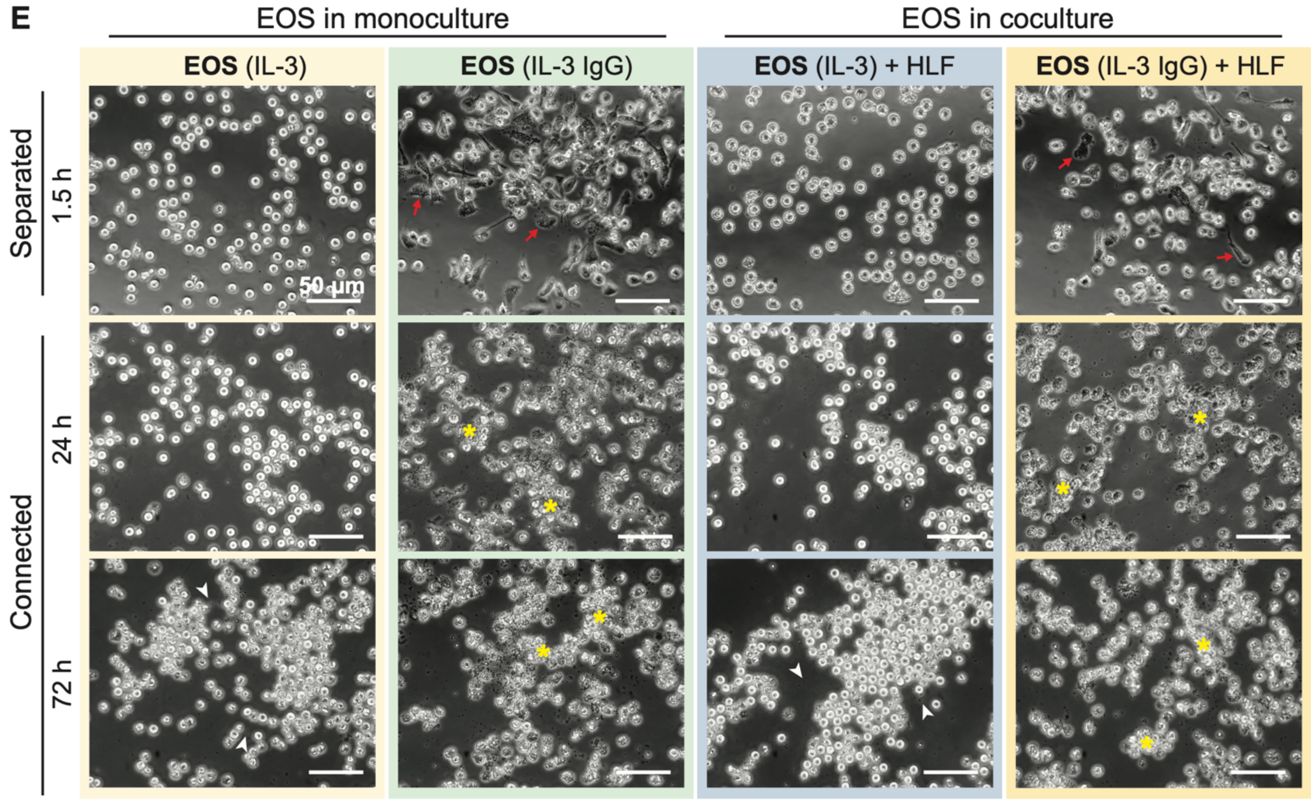


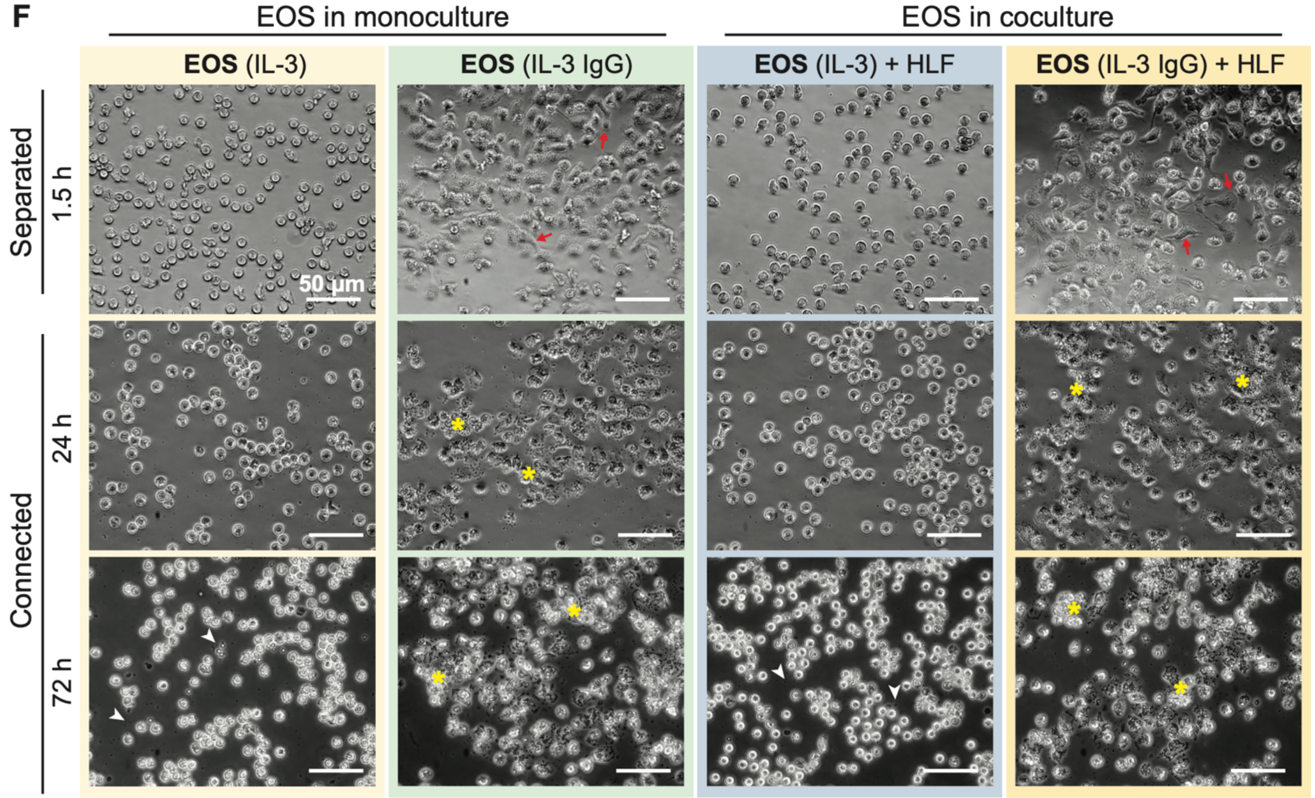


(continued next page)

**Supplementary Figure 4.** Phase contrast images of eosinophils activated with IL-3 ("IL-3") and in some conditions also seeded onto HA-IgG ("IL-3 IgG") in coculture devices at the indicated times. Representative images from 6 donor pairs are shown: **(A)** eosinophil donor 1 and HLF donor 1, **(B)** eosinophil donor 1 and HLF donor 2, **(C)** eosinophil donor 2 and HLF donor 1, **(D)** eosinophil donor 2 and HLF donor 2, **(E)** eosinophil donor 3 and HLF donor 1, **(F)** eosinophil donor 3 and HLF donor 2. The chambers remained separated for 1.5 h and then were connected for up to 72 h. Red arrows show eosinophils spreading on HA-IgG. Yellow asterisks show aggregates of eosinophil cell debris. White arrowheads show eosinophil cell-free granules. Images are representative of 4 culture replicates.

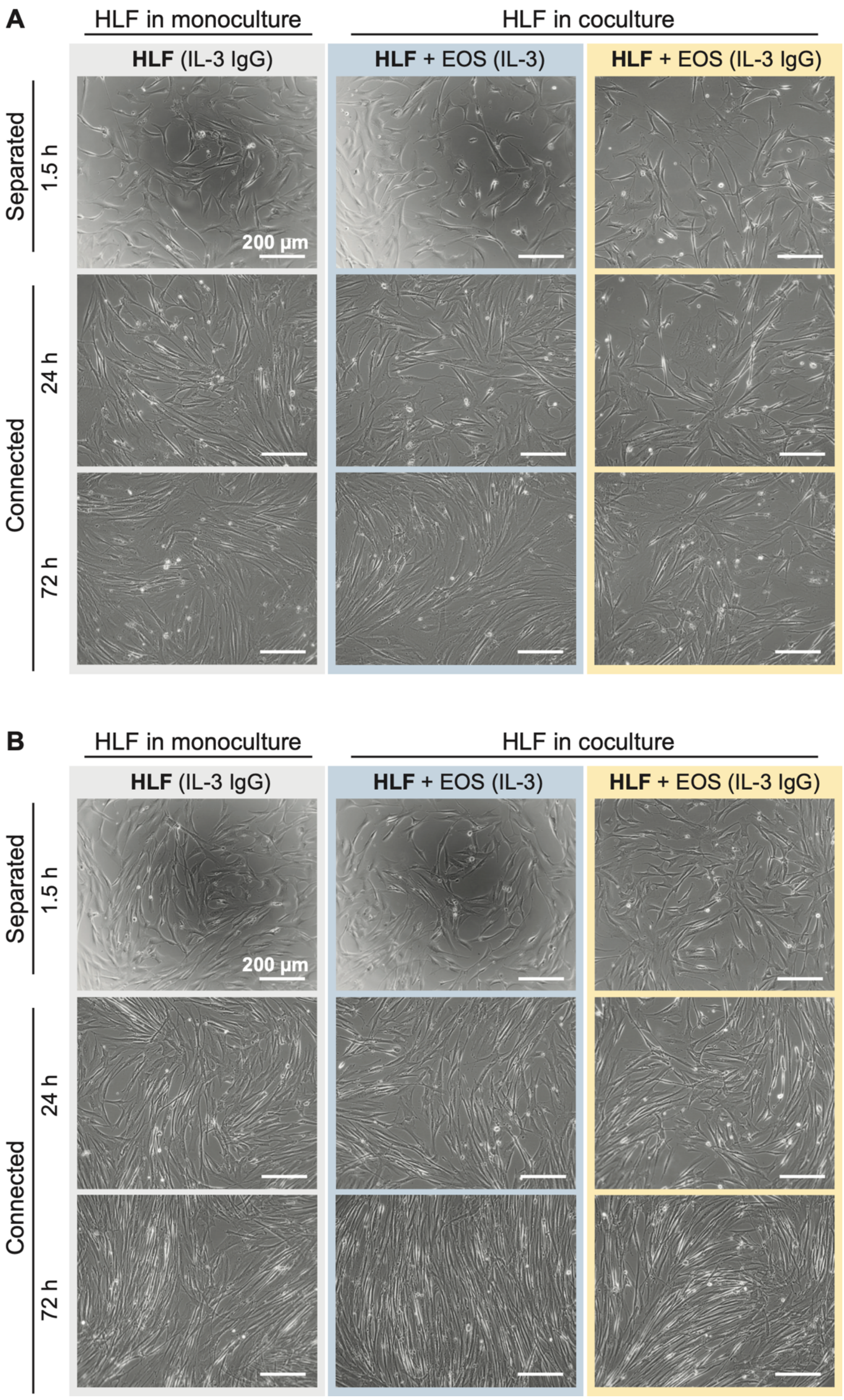


(continued next page)

**Supplementary Figure 5.** Phase contrast images of HLFs in coculture devices at the indicated times. Representative images from 2 donor pairs are shown: **(A)** eosinophil donor 2 and HLF donor 1, **(B)** eosinophil donor 2 and HLF donor 2. The chambers remained separated for 1.5 h and then were connected for up to 72 h. Images are representative of 4 culture replicates, as well as 3 eosinophil donors and 2 HLF donors.

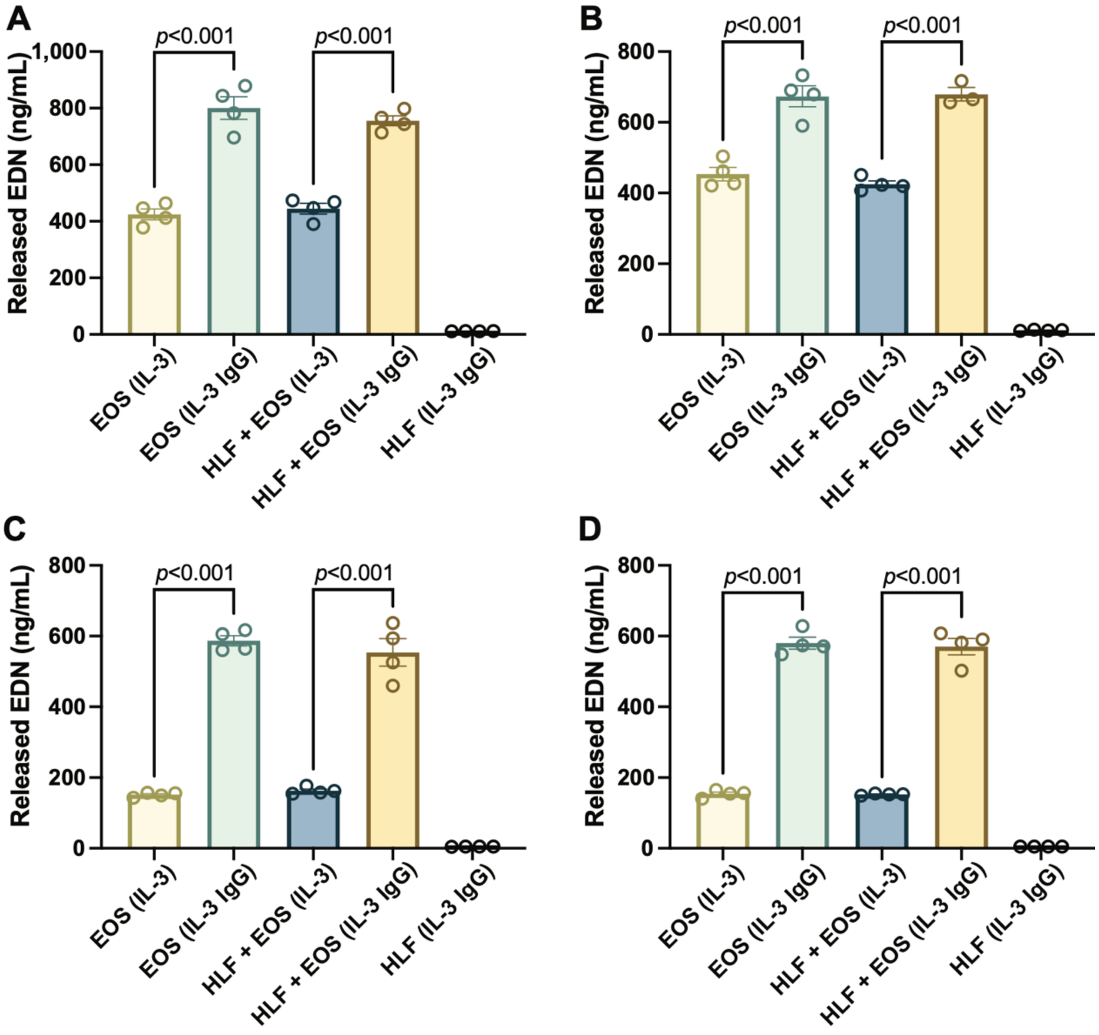


**Supplementary Figure 6.** Levels of released eosinophil-derived neurotoxin (EDN) in culture media after 24 h measured by ELISA. Eosinophils release the highest level of EDN under IL-3 IgG conditions regardless of coculture or monoculture. Results from 4 donor pairs are shown: **(A)** eosinophil donor 1 and HLF donor 1, **(B)** eosinophil donor 1 and HLF donor 2, **(C)** eosinophil donor 3 and HLF donor 1, **(D)** eosinophil donor 3 and HLF donor 2. The 2 donor pairs with eosinophil donor 2 were excluded due to protocol differences in ELISA. Data are expressed as mean ± SEM (n=4); each point is one culture replicate. Data were analyzed using one-way ANOVA, followed by Holm-Šídák's multiple comparisons test.

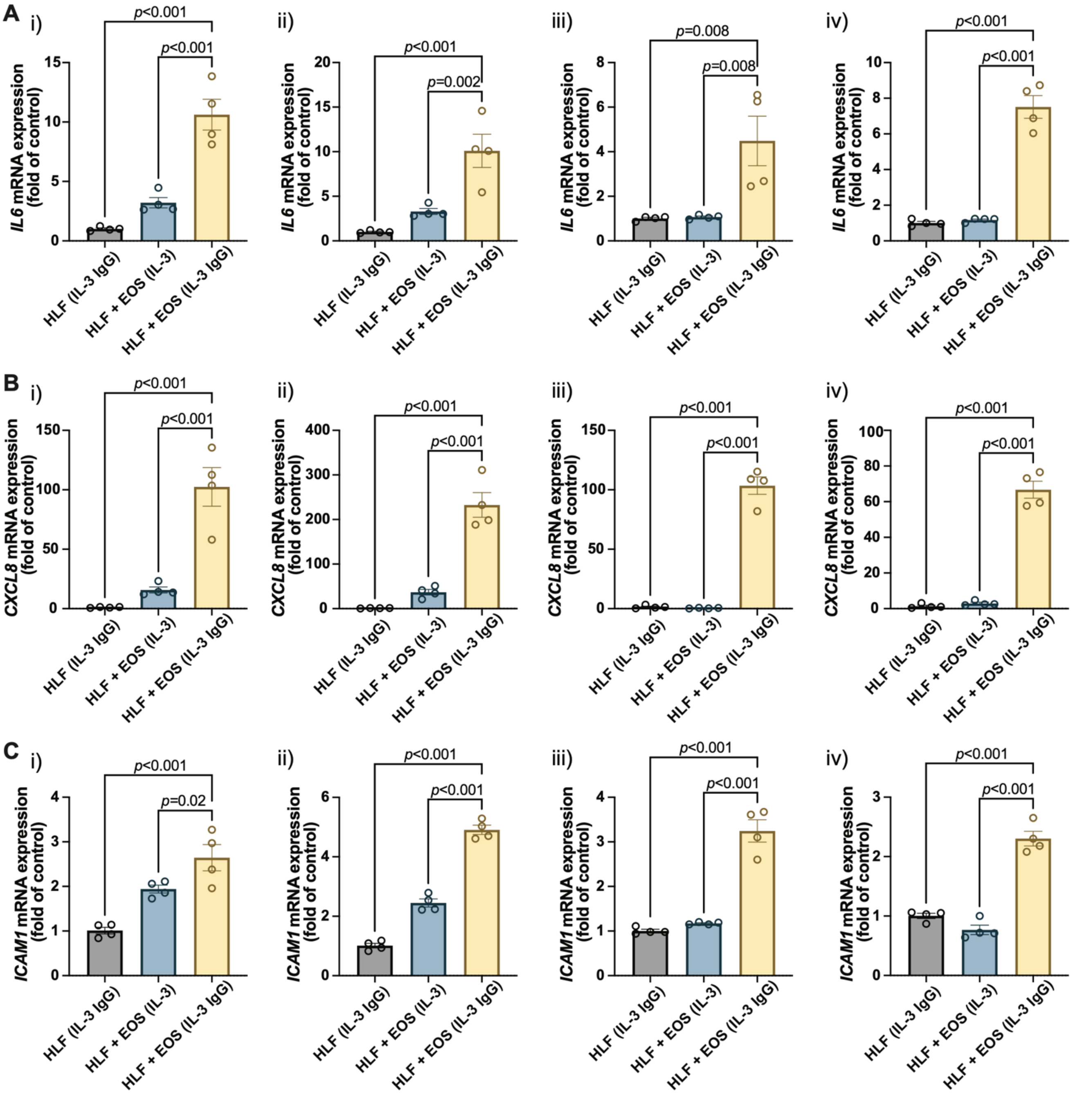


**Supplementary Figure 7.** Proinflammatory genes are upregulated in HLFs cocultured with degranulating eosinophils. HLFs in coculture with degranulating eosinophils ("HLF + EOS (IL-3 IgG)") for 72 h showed the highest mRNA level upregulation of **(A)** *IL6*, **(B)** *CXCL8*, and **(C)** *ICAM1*. Results from 4 donor pairs are shown: **(i)** eosinophil donor 1 and HLF donor 1, **(ii)** eosinophil donor 1 and HLF donor 2, **(iii)** eosinophil donor 2 and HLF donor 1, **(iv)** eosinophil donor 2 and HLF donor 2. Data from the 2 donor pairs with eosinophil donor 3 were not included because the samples were used to develop a separate readout. Data are expressed as mean ± SEM (n=4); each point is one culture replicate. Data were analyzed using one-way ANOVA, followed by Holm-Šídák's multiple comparisons test.

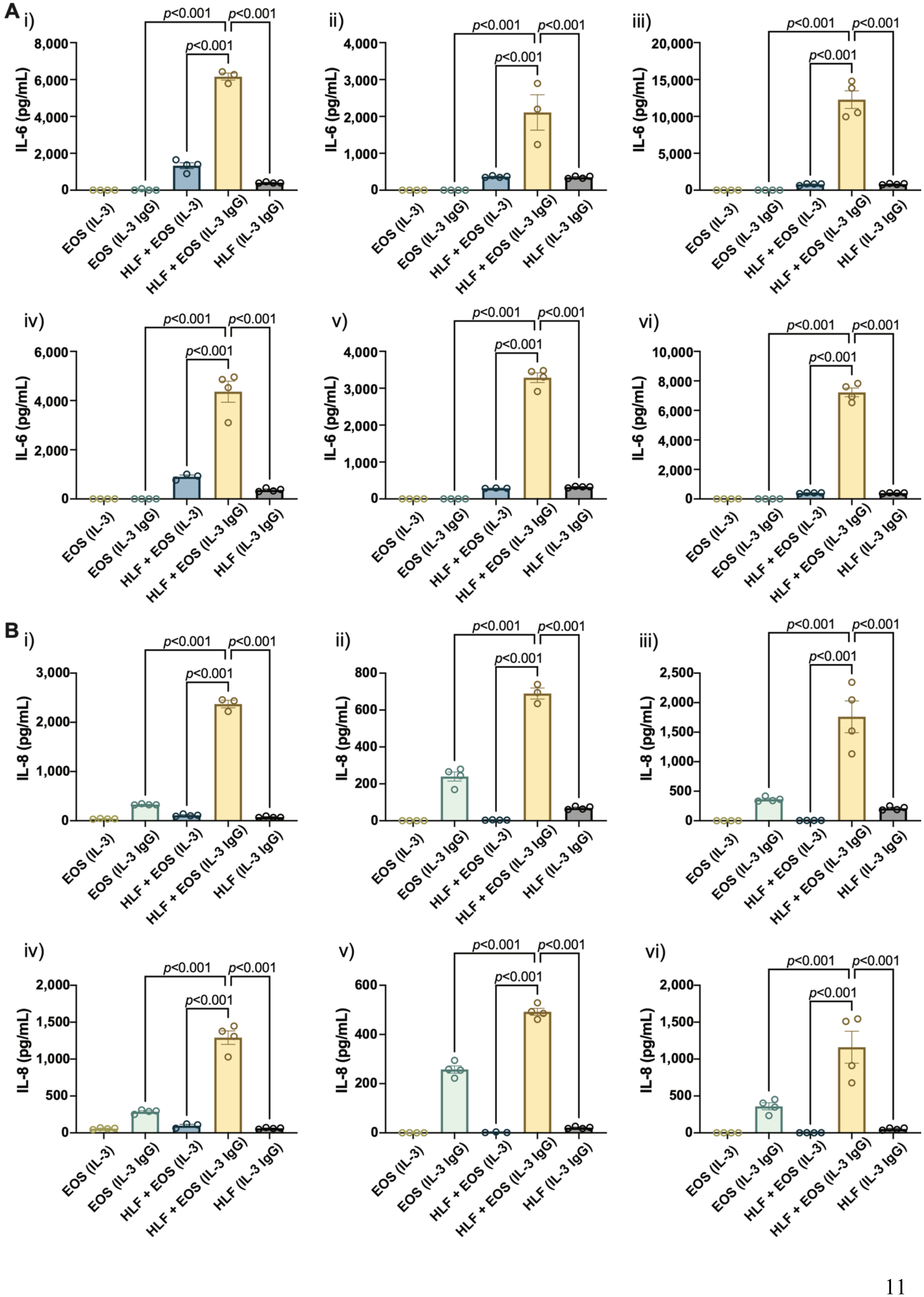
A
B
i)
ii)
iii)
iv)
v)
vi)
IL-6 (pg/mL)
IL-8 (pg/mL)
EOS (IL-3)
EOS (IL-3 IgG)
HLF + EOS (IL-3)
HLF + EOS (IL-3 IgG)
HLF (IL-3 IgG)
p<0.001

**Supplementary Figure 8.** The coculture of HLFs with degranulating eosinophils releases the highest levels of proinflammatory cytokines compared to HLF and eosinophil monocultures and other control conditions. Levels of soluble factors, **(A)** IL-6 and **(B)** IL-8, were measured in the conditioned media after 72 h in culture. Results from 6 donor pairs are shown: **(i)** eosinophil donor 1 and HLF donor 1, **(ii)** eosinophil donor 2 and HLF donor 1, **(iii)** eosinophil donor 3 and HLF donor 1, **(iv)** eosinophil donor 1 and HLF donor 2, **(v)** eosinophil donor 2 and HLF donor 2, **(vi)** eosinophil donor 3 and HLF donor 2. Data are expressed as mean ± SEM (n=3-4); each point is one culture replicate. Data were analyzed using one-way ANOVA, followed by Holm-Šídák's multiple comparisons test.

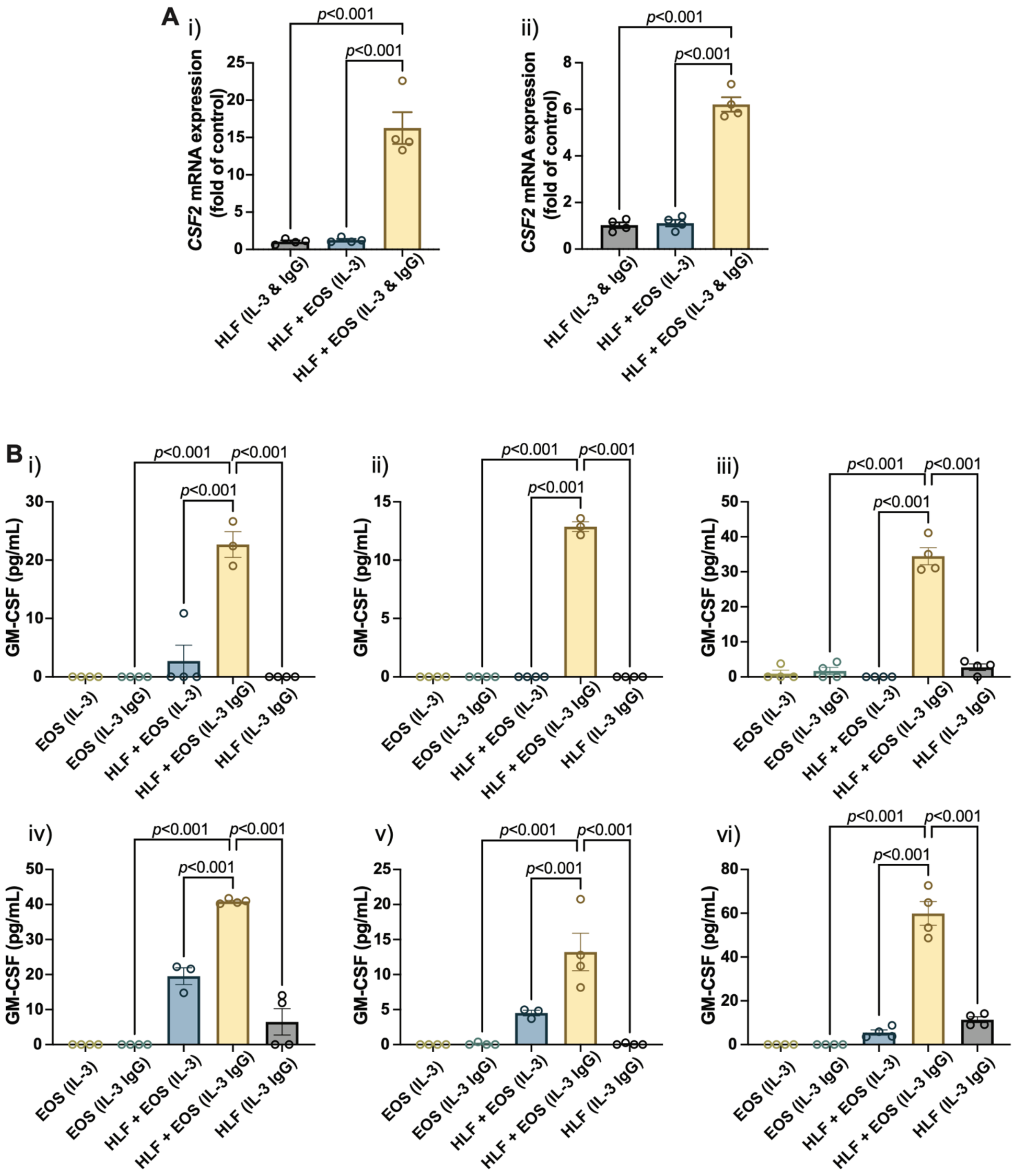


**Supplementary Figure 9.** The coculture of HLFs with degranulating eosinophils induces the highest levels of expression and secretion of *CSF2*/GM-CSF than other control conditions. **(A)** mRNA levels of *CSF2* in HLFs after 72 h in culture. Results from 2 donor pairs are shown: **(i)** eosinophil donor 2 and HLF donor 1, **(ii)** eosinophil donor 2 and HLF donor 2. Data from the 4 donor pairs with eosinophil donors 1 and 3 were not included, because the samples were limited and used up to develop a separate readout. **(B)** Levels of GM-CSF measured in the conditioned media after 72 h in culture. Results from 6 donor pairs are shown: **(i)** eosinophil donor 1 and HLF donor 1, **(ii)**

eosinophil donor 2 and HLF donor 1, **(iii)** eosinophil donor 3 and HLF donor 1, **(iv)** eosinophil donor 1 and HLF donor 2, **(v)** eosinophil donor 2 and HLF donor 2, **(vi)** eosinophil donor 3 and HLF donor 2. Data are expressed as mean ± SEM (n=3-4); each point is one culture replicate. Data were analyzed using one-way ANOVA, followed by Holm-Šídák's multiple comparisons test.

#### 2 Supplementary Tables

**Supplementary Table 1.** Calculation of eosinophil seeding densities in 96-well plates (Esnault et al., 2017) or outer chambers of the microfluidic coculture device. The surface area seeding density was kept consistent in both cultureware.

| | **96-well plate** | **Device outer chamber** |
|---|---|---|
| Media volume | 100 µL | 36 µL |
| Seeding area | 32 $mm^2$ | 29.3 $mm^2$ |
| Volume density | $1 \times 10^6$ cells/mL | $2.6 \times 10^6$ cells/mL |
| Area density | 3125 cells/$mm^2$ | |
| Total cell number per well | $1 \times 10^5$ cells | $9.2 \times 10^4$ cells |

**Supplementary Table 2.** Forward and reserve sequences of primers used in reverse transcription quantitative polymerase chain reaction (RT-qPCR). *GUSB*: glucuronidase β. *IL6*: interleukin 6. *CXCL8*: C-X-C motif chemokine ligand 8. *ICAM1*: intercellular adhesion molecule 1. *CSF2*: colony stimulating factor 2.

| **Primer** | **Forward** | **Reverse** |
|---|---|---|
| *GUSB* | CAGGACCTGCGCACAAGAG | AGCGTGTCGACCCCATTC |
| *IL6* | TGCAGATGAGTACAAAAGTCCTGAT | GTGGTTATTGCATCTAGATTCTTTGC |
| *CXCL8* | CTTGGCAGCCTTCCTGATTT | TTCTTTAGCACTCCTTGGCAAAA |
| *ICAM1* | GCCAGGAGACACTGCAGACA | TGGCTTCGTCAGAATCACGTT |
| *CSF2* | TGAATGAAACAGTAGAAGTCATCTCAGAAAT | GCTCCAGGCGGGTCTGTA |
